# Paracrine Activin-A signaling promotes melanoma growth and metastasis through immune evasion

**DOI:** 10.1101/144857

**Authors:** Prudence Donovan, Olivier A. Dubey, Susanna Kallioinen, Katherine W. Rogers, Katja Muehlethaler, Patrick Müller, Donata Rimoldi, Daniel B. Constam

**Affiliations:** Ecole Polytechnique Fédérale de Lausanne (EPFL) SV ISREC, Station 19, CH-1015 Lausanne, Switzerland; Centre Ludwig de 1’UNIL pour la recherche sur le cancer, Ch. des Boveresses 155, 1066 Epalinges, Switzerland; Friedrich Miescher Laboratory of the Max Planck Society, Spemannstraße 39, 72076 Tübingen, Germany

## Abstract

The secreted growth factor Activin-A of the TGFβ family and its receptors can promote or inhibit several cancer hallmarks including tumor cell proliferation and differentiation, vascularization, lymphangiogenesis and inflammation. However, a role in immune evasion and its relationship with tumor-induced muscle wasting and tumor vascularization, and the relative contributions of autocrine versus paracrine Activin signaling remain to be evaluated. To address this, we compared the effects of truncated soluble Activin receptor II B as a ligand trap, or constitutively active mutant type IB receptor versus secreted Activin-A or the related ligand Nodal in mouse and human melanoma cell lines and tumor grafts. We found that while cell-autonomous receptor activation arrested tumor cell proliferation, Activin-A secretion stimulated melanoma cell dedifferentiation and tumor vascularization by functional blood vessels, and it increased primary and metastatic tumor burden and muscle wasting. Importantly, in mice with impaired adaptive immunity, the tumor-promoting effect of Activin-A was lost despite sustained vascularization and cachexia, suggesting that Activin-A promotes melanoma progression by inhibiting anti-tumor immunity. Paracrine Activin-A signaling emerges as a potential target for personalized therapies, both to reduce cachexia and to enhance the efficacy of immunotherapies.

## INTRODUCTION

In melanoma research, targeted inhibitors and immune checkpoint therapies are available but their efficacies remain limited by acquired and intrinsic drug resistance mechanisms (Lau et al., 2016). Patients with resistant melanoma have a poor prognosis due to high probability of metastasis, correlating with progression from superficial spreading to invasive vertical growth. Tumor-initiating potential and metastasis also correlate with low immunogenic profiles of subpopulations of cells and phenotypic plasticity marked by pseudo-epithelial-to-mesenchymal transitions (EMT) and the ability to reversibly switch between proliferative and invasive gene signatures (Caramel et al., 2013, Hoek et al., 2008, Schatton and Frank, 2009, Widmer et al., 2012). Thus, elucidating mechanisms of tumor immune evasion and their coupling to cancer cell plasticity is critical to develop effective immunotherapies.

Known local cues in the tumor microenvironment that regulate melanoma cell plasticity and anti-tumor immunity include transforming growth factor beta (TGFβ). In many tumors including melanoma, TGFβ facilitates or inhibits tumor progression, depending on the context (Bellomo et al., 2016, Perrot et al., 2013). In normal melanocytes, TGFβ induces cell cycle arrest and apoptosis (Alanko and Saksela, 2000, Rodeck et al., 1991), but this response is attenuated in cells from benign nevi despite sustained activation of downstream SMAD2,3 transcription factors (Rodeck et al., 1999). SMAD-binding sites in the *PAX3* gene mediating repression of pigment synthesis likely contribute to melanoma cell plasticity and phenotype switching (Pinner et al., 2009, Yang et al., 2008). TGFβ immunostaining was found to correlate with the invasive vertical growth phase and metastasis (Van Belle et al., 1996), and blockade of downstream SMAD2,3 transcription factors by overexpression of antagonistic SMAD7 in 1205Lu melanoma cells inhibited tumorigenicity and bone metastasis (Javelaud et al., 2005, Javelaud et al., 2007). In metastatic B16F10 mouse melanoma, TGFβ also reduces Natural Killer cell-mediated tumor rejection while promoting the differentiation of immunosuppressive Foxp3+ regulatory T cells in tumor beds and draining lymph nodes (Chen et al., 2003, Gorelik and Flavell, 2001, Turk et al., 2004). However, the functionally relevant active TGFβ in this model is provided by immature myeloid dendritic cells in draining lymph nodes, likely because B16F10 and other melanoma cells themselves secrete TGFβ only in latent form (Ghiringhelli et al., 2005, Yin et al., 2012).

Melanoma cell lines and tumors also frequently express Activin-A (Heinz et al., 2015, Hoek et al., 2006). Activin-A stimulates the same SMAD transcription factors as TGFβ, even though it derives from distinct precursor dimers encoded by the *Inhibin βA* (*INHβA*) gene and binds distinct complexes of the Activin/Nodal type I and II receptors Acvr1b/ALK4 or Acvr1c/ALK7 and Acvr2 (reviewed in Hedger et al., 2011, Sozzani and Musso, 2011, Walton et al., 2011). Activin-A can mediate protective or tumorigenic effects (reviewed in Loomans and Andl, 2014). For example in mouse models of pancreatic cancer, Activin-A promotes tumor progression by inhibiting cell differentiation (Lonardo et al., 2011, Togashi et al., 2015). Paradoxically, its receptor ALK4 (ACVR1B) primarily mediates tumor suppressive functions and is frequently deleted in clinical samples (Qiu et al., 2016, Togashi et al., 2014). Cytostatic and proapoptotic signaling by Activin-A has also been reported in human melanoma cell lines, although this activity is counteracted by the secreted antagonist Follistatin (FST-1) (Stove et al., 2004). Immunohistochemical analysis detected Activin-A staining in superficially spreading melanoma, whereas benign nevi and metastatic lesions showed elevated expression of Follistatin (Heinz et al., 2015). Interestingly, however, gain of transgenic Activin-A expression in the A375 human melanoma xenograft model did not alter tumor growth or metastasis, even though it reduced tumor lymphangiogenesis, a risk factor of poor prognosis (Heinz et al., 2015). Also in xenograft models of other cancers, overexpression of a dominant negative mutant receptor revealed no adverse functions for Activin-A or related ligands, with the exception of tumor-induced systemic weight loss (cachexia) through remote Smad2,3 activation in skeletal muscles (Li et al., 2007, Zhou et al., 2010).

Here, we asked whether a potential tumorigenic role for Activin signaling in melanoma may be curtailed in xenografts by the absence of a functional immune system. To address this, we monitored the expression of *INHβA* in public datasets and in a panel of melanoma tumors and cell lines, and we performed loss- and gain-of-function studies, respectively, in human xenografts and in the moderately metastatic syngeneic B16F1 mouse melanoma model using dominant negative or ligand-independent mutant Activin/Nodal receptors or Activin-A lentiviral transgenes. Our comparison of melanoma grafts in tumor hosts of different genetic backgrounds reveals a novel protumorigenic and prometastatic function specifically for paracrine Activin-A signaling acting on adaptive immunity, and that this function can be uncoupled from immune-independent anorexic and proangiogenic effects and from autocrine effects on melanoma cell differentiation. Our findings suggest that gain of Activin-A expression in human melanoma should be considered as a potential new target for improved immunotherapies.

## RESULTS

### Human melanoma and other skin cancers frequently express Activin-A rather than NODAL

To assess potential roles of *INHβA* in melanoma, we queried a gene expression profiling dataset of 42 primary cutaneous tumors (Riker et al., 2008). We found that *INHβA* expression was significantly elevated both in primary cutaneous cancers and in metastatic melanoma, whereas the antagonist Follistatin tended to decrease in melanoma *in situ* (n=2, not significant) compared to normal skin. In contrast to *INHβA*, other Activin receptor ligands encoded by *NODAL* or *GDF3* were neither upregulated in this dataset nor in a TCGA collection of 474 skin cutaneous melanoma (**Figs. 1A, S1**). Elevated *INHβA* expression significantly clustered with a standard gene expression signature of invasive rather than proliferative melanoma cell lines as determined by Heuristic Online Phenotype Prediction (HOPP) analysis (**Fig. 1B**) (Widmer et al., 2012). Furthermore, while RT-PCR analysis confirmed *INHβA* expression in more than half of a panel of 20 human melanoma cell lines, the related ligands encoded by *INHβB* or *NODAL* were rarely transcribed or undetectable, respectively (**Figs. 1C, S2**). Although survival did not correlate with either *INHβA* or *NODAL*, low expression of the Activin antagonist *Follistatin* in the TCGA dataset was associated with worse outcome (**Fig. S3**). These results are consistent with a potential role for Activin-A signaling in melanoma progression.

**Figure 1.**
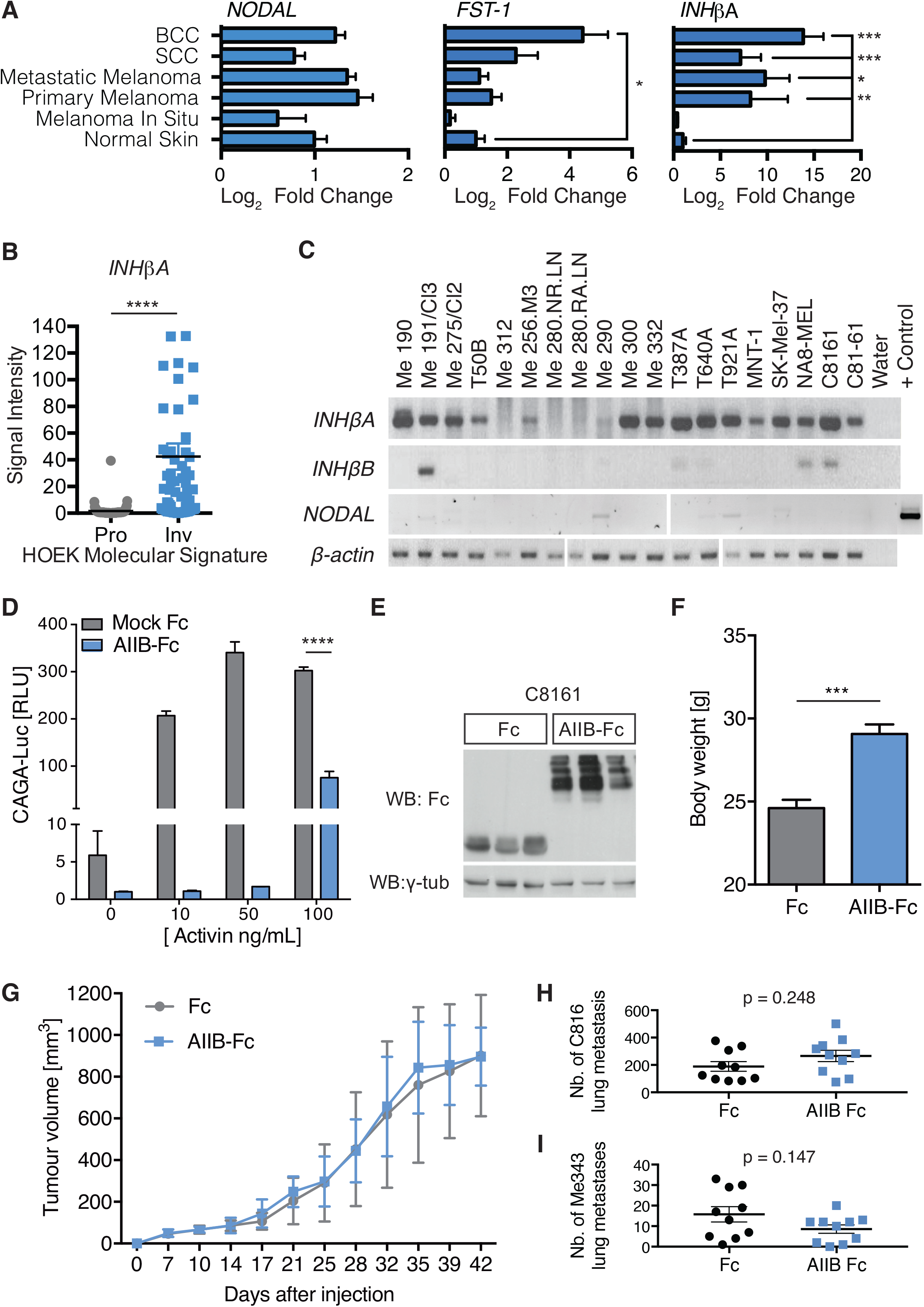
Cachexia induced by endogenous Activin-A in human melanoma xenografts does not promote tumor progression. A) Expression of *NODAL*, Follistatin (*FST-1*) and *INHβA* (encoding Activin-A) mRNAs in normal skin compared to 42 primary cutaneous tumors comprising 14 melanoma, 11 squamous cell, 15 basal cell skin cancers and 40 melanoma metastases (Riker et al., 2008). *p < 0.05; **p < 0.01; ***p < 0.001 B) Relative *INHβA* mRNA levels in 536 human melanoma cell lines distinguished by proliferative (Pro) or invasive (Inv) gene expression signatures (Widmer et al., 2012). C) RT-PCR analysis of the indicated mRNAs in total RNA from human melanoma cell lines or fetal brain (+Control). D) Normalized expression of the Smad3 luciferase reporter CAGA-Luc in HEK293T cells treated with the indicated concentration of Activin-A or with conditioned media of C8161 melanoma cells expressing lentiviral AIIB-Fc or Fc alone (mock). Data show the average fold change of 3 experiments ± SD, ****p< 0.0001 at all concentrations. E) Western blot analysis of lentiviral AIIB-Fc or Fc alone in intradermal C8161 melanoma xenograft tumors, and F) body weights of Foxn1nu/nu hosts at the time of tumor resection (***p < 0.01). G) Growth curves of C8161 melanoma were unaffected by AIIB-Fc expression. H) Number of superficial lung metastases of C8161-Fc (MockFc) or C8161-AIIBFc melanoma cells 4 weeks after injection into the tail vein of Foxn1^nu/nu^ mice (n=10 per group, p=0.25, two-tailed Mann-Whitney). I) As in H), but using human Me343 melanoma cells (n=10 per group, p=0.14)

### Inhibition of endogenous ActRIIB ligands in human C8161 melanoma cells inhibits cachexia but not tumor progression in immunocompromised nude mice

Previous analysis of Activin-A functions in tumor xenograft models uncovered potent anorexic activity but no important roles in tumor progression (Heinz et al., 2015, Zhou et al., 2010). Instead, increased aggressiveness in human melanoma has been attributed to a secreted Nodal-like activity that was initially detected by injecting C8161 melanoma cells into zebrafish embryos (Topczewska et al., 2006). Since we observed no *NODAL* expression in human melanoma, we decided to reassess how C8161 cells stimulate Nodal/Activin receptors in zebrafish. To address this, we grafted C8161 cells into wild-type or mutant zebrafish embryos lacking maternal and zygotic expression of the Nodal co-receptor *one-eyed pinhead* (MZ*oep*) (Gritsman et al., 1999). To validate that human NODAL signaling requires Oep in zebrafish, we injected 50 pg of mRNA encoding human Nodal at the one-cell stage and monitored induction of the Smad2,3 target gene *gsc* during early gastrulation. We found that both human Nodal and the zebrafish homolog Squint were active in wild-type but not in MZ*oep*, whereas human Activin-A induced *gsc* independently of Oep (**Fig. S4A**). Moreover, both wild-type and MZ*oep* embryos ectopically upregulated *gsc* as well as *ntl* around grafted C8161 melanoma cells (**Fig. S4B**). These results strongly argue for secreted Activin-A and against Nodal as the mediator of Smad2,3 signaling secreted by C8161 melanoma grafts in zebrafish.

To test whether melanoma may grow faster when forced to express Nodal, C8161 cells transduced with Nodal lentivirus (**Fig. S2D**) were grafted subcutaneously into FoxN1^nu/nu^ mice. C8161 xenograft tumor growth was not increased by Nodal compared to cells transduced with empty vector, indicating that it was not limited by lack of Nodal expression (**Fig. S4E, F**). To assess the function of endogenous ligands in melanoma xenografts, we transduced human C8161 cells with lentivirus expressing an Fc fusion of Activin type IIB receptor extracellular domain (AIIB-Fc) or Fc alone. Lentivirally transduced AIIB-Fc inhibited endogenous Activin activity in conditioned medium of C8161 cells *in vitro* and was readily detected by immunoblotting in C8161 tumor xenografts (**Fig. 1D, E**). As expected, expression of AIIB-Fc in C8161 xenografts significantly protected FoxN1^nu/nu^ hosts against loss of body weight and muscle mass (**Fig. 1F**). However, despite potent systemic inhibition of cachexia, soluble AIIB-Fc receptor neither diminished intradermal tumor growth nor experimental lung metastases after tail vein injection (**Fig. 1G, H**). Additionally, secondary tumor outgrowths following resection of the primary C8161 graft and spontaneous lung or lymph node metastases were not significantly inhibited (**Fig. S5A-C**), nor did AIIB-Fc transduction inhibit experimental lung metastases of Me343 cells which also express *INHβA* (**Figs. 1I, S2G**). These results suggest that C8161 xenografts grow and metastasize independently of Activin signaling and of associated systemic cachexia, possibly because of the lack of functional adaptive immunity.

### Activin-A gain of function promotes phenotype switching of mouse melanoma cells and tumor progression

To evaluate potential tumorigenic or prometastatic effects of Activin-A in immunocompetent hosts, we transduced *INHβA* in the moderately metastatic syngeneic B16F1 mouse melanoma model (Fidler, 1973) using lentiviruses for GFP alone (CTRL) or GFP together with Activin-A (INHβA). Western blot analysis of B16F1-conditioned medium and cell lysates confirmed Activin-A secretion which increased the accumulation of pSmad2 compared to CTRL cells that do not secrete Activin-A (**Fig. 2A**). Transfection of the SMAD3 luciferase reporter CAGA-Luc showed elevated autocrine Activin-A signaling in *INHβA*-compared to CTRL-transduced B16F1 cells, and this increase was blocked upon treatment with Follistatin (**Fig. 2B**, left panel). Conditioned medium from INHβA-transduced B16F1 cells also stimulated the expression of transfected CAGA-Luc in HEK293T reporter cells (**Fig. 2B**, right panel). Furthermore, gain of Activin-A expression in B16F1 cells stimulated their migration in a scratch wound assay and markedly reduced their melanin content without altering their cell proliferation or viability (**Fig. 2C-E**), consistent with a potential autocrine function in promoting a phenotypic switch.

**Figure 2.**
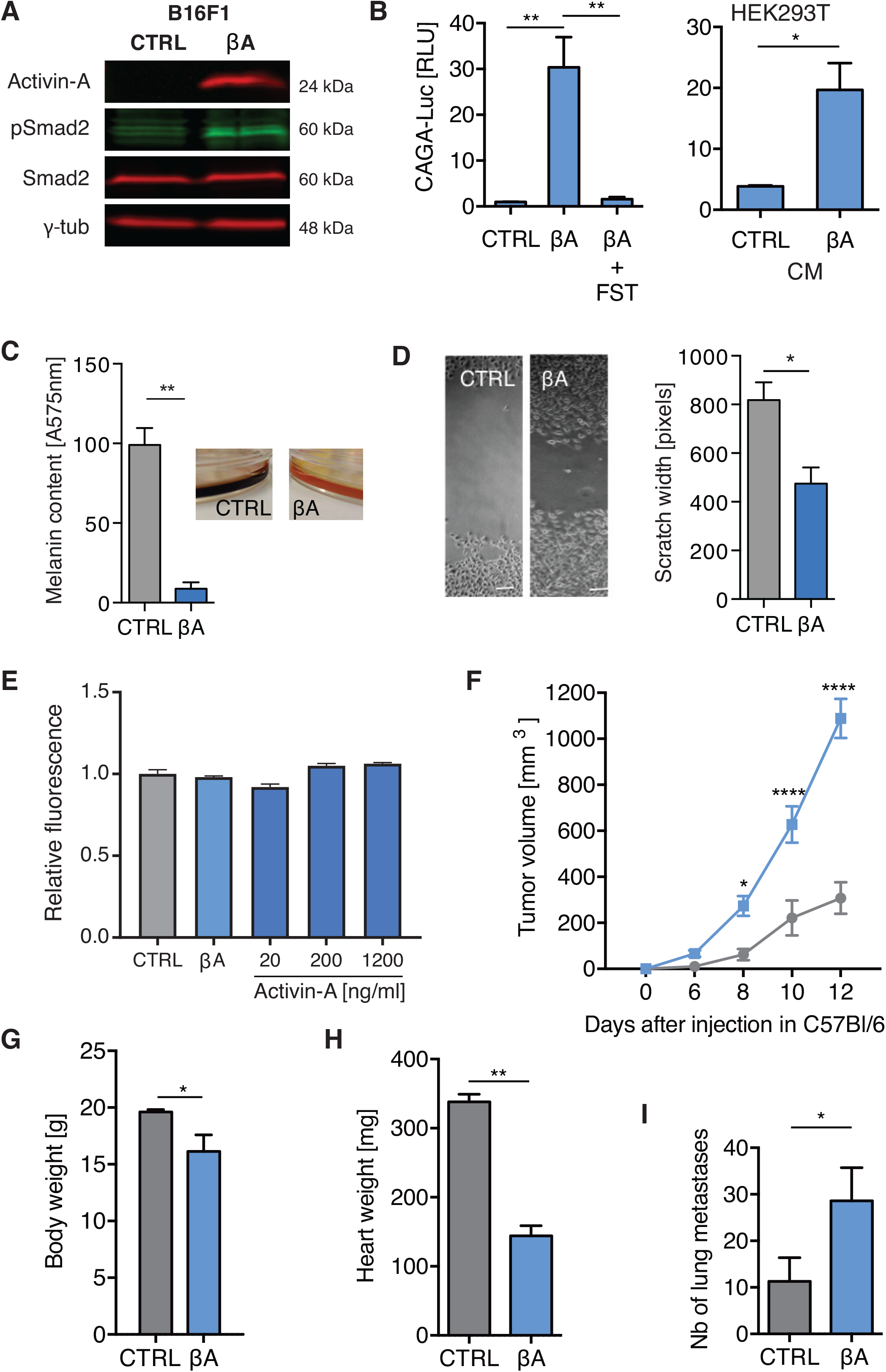
Gain of Activin-A expression stimulates phenotype switching and syngeneic tumor growth in B16F1 mouse melanoma. A-B) Western blot of A) secreted Activin-A and B) CAGA-Luc expression in B16F1:GFP cells (left panel) transduced with *INHβA* (βA) or control lentivirus (CTRL). Where indicated, cells were treated with Follistatin (FST, 300 ng/ml). CAGA-Luc expression in HEK293T cells treated with conditioned media of B16:GFP.ßA or B16:GFP.CTRL cells is also shown (right panel in B). Data represent mean±SEM of three experiments; * p<0.05, ** p<0.01. C) Light absorbance by pigment in medium of transduced B16F1:GFP cells (mean±SEM, n=3; **p<0.01). D) Representative micrographs and widths of scratch wounds 20 hours after wounding of cell monolayers (mean±SEM from 3 experiments, *p<0.05). E) Alamar blue quantification of live B16F1 cells transduced with control or *INHβA* lentivirus, or treated with the indicated concentrations of Activin-A for 48 hours (mean±SEM, n=3, no significant differences). F-H) Growth curves of B16F1:GFP syngeneic intradermal grafts with Activin-A (βA) or without (CTRL) in C57BL/6 mice, and body (G) and heart (H) weights at the time of tumor resection. Data represent mean±SEM of 20 animals/group; *p<0.05, **p<0.01, ****p<0.0001. I) Lung metastases of tail-vein injected B16F1:GFP βA or CTRL cells (n=15 each). Data represent mean±SEM of 3 experiments; *p<0.05.

To determine if oncogenic effects of Activin-A gain-of-function may promote tumor growth *in vivo*, B16F1 grafts were intradermally inoculated on the right flank of 8-10 week old female C57BL/6 mice. Mice injected with INHβA-transduced cells grew significantly larger tumors, combined with loss of body weight and cardiac muscle wasting (**Fig. 2F-H**). Compared to vector control, *INHβA* also increased the number of experimental lung metastases formed by tumor cells that were injected into the tail vein (**Fig. 2I**). Taken together, these results show that Activin-A signaling can promote melanoma growth and metastasis in immunocompetent hosts.

### Sustained cell autonomous ALK4 signaling inhibits B16F1 tumorigenesis rather than stimulating it

To validate whether the tumorigenic function of Activin-A involves cell non-autonomous paracrine signaling, we transduced B16F1 cells with lentivirus expressing a constitutively active ligand-independent truncated form of ALK4 (caALK4). Induction of caALK4 by doxycycline led to increased Smad2,3 phosphorylation and expression of the SMAD3-dependent luciferase reporter CAGA-Luc (**Fig. 3A-C**) and reduced pigment secretion (**Fig. S6A**). However, caALK4 expression decreased below detection within four subsequent cell passages despite continuous presence of doxycycline (**Fig. 3A**). EdU incorporation, AnnexinV and propidium iodide flow cytometry and cleaved Caspase3 immunofluorescence staining after up to 3 days of caALK4 induction *in vitro* revealed no overt cell cycle inhibition or apoptosis (**Fig. S7A-D** and data not shown), even though the number of cells detected by Alamar blue assays after 3 days of caALK4 induction *in vitro* was reduced (**Fig. 3D**), indicating impaired cell viability. A similar reduction in the number of viable cells was induced by ligand-independent caALK4 in human C8161 melanoma cells. To investigate the impact on tumor growth, B16F1 cells transduced with caALK4 were grafted intradermally into syngeneic C57BL/6 hosts, followed by treatment with doxycycline or empty vehicle until the endpoint of the experiment. Tumors induced no overt loss of body weight (**Fig. 3E**), and they grew significantly less in doxycycline-fed animals than in vehicle-treated controls (**Fig. 3F**), indicating that sustained autocrine Activin receptor signaling does not stimulate but rather suppresses tumor growth *in vivo.* Immunostaining at the endpoint of the experiment revealed increased apoptosis marked by cleaved Caspase-3 (**Figs. 3G, S7E**) while Ki67-stained proliferating cells appeared to decrease, albeit not significantly (**Fig. 3H-I** and **Supp. Fig. S7F**). We conclude that while paracrine functions of Activin-A promote cachexia and tumor growth, sustained autocrine receptor signaling *in vivo* reduces B16F1 cell survival and tumorigenicity.

**Figure 3.**
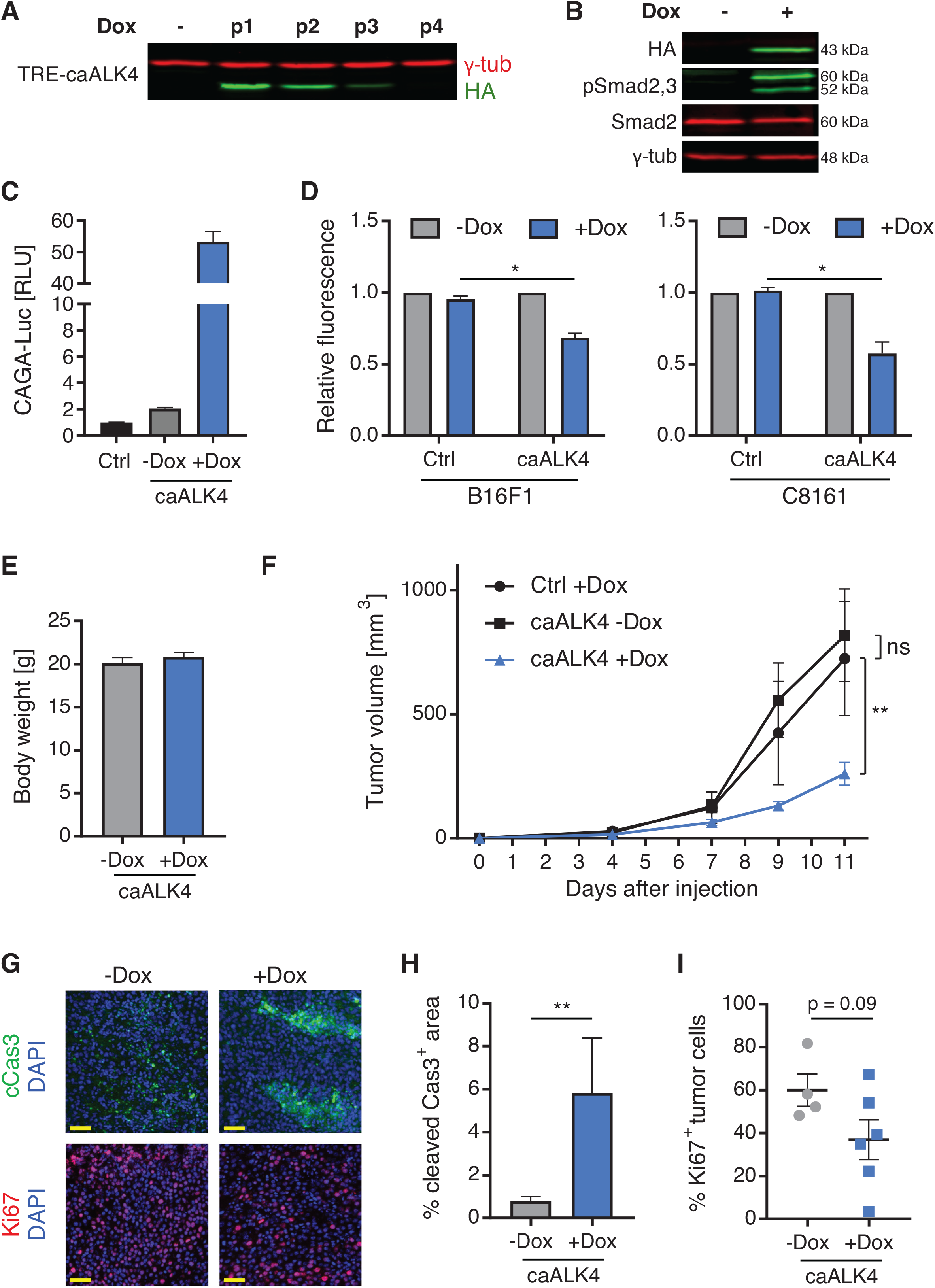
Sustained autocrine Activin receptor signaling inhibits B16F1 cell survival and tumorigenesis rather than stimulating it. A, B) anti-HA and phospho-Smad2,3 Western blot of B16F1 cells treated with doxycycline during 4 cell passages (A) or 24 hours (B) after lentiviral infection of inducible HA-tagged caALK4 (rtTA-caALK4). C) CAGA-Luc expression in B16F1 (Ctrl) or B16F1 rtTA-caALK4 cells ± doxycycline for 24h. Data represent mean±SEM of 3 experiments. D) Alamar blue assay of B16F1 or C8161 (Ctrl) and B16F1 rtTA-caALK4 or C8161 rtTA-caALK4 cells cultured for 3 days in medium with or without doxycycline. Data represent mean±SEM of 2 experiments, *p<0.05. E) Body weights of C57BL/6 hosts at the time of tumor resection after feeding with or without doxycycline, and F) growth curves of intradermal syngeneic grafts of B16F1 rtTA (Ctrl, n=15) and B16F1 rtTA-caALK4 cells (n=15). Data represent mean±SEM of 3 experiments, **p<0.01. G) Cleaved Caspase-3 (cCas3) and Ki67 immunofluorescence staining of syngeneic B16F1 tumors expressing caALK4 (+Dox) or not (−Dox). Scale bar: 50μm. Corresponding whole-section images are shown in Fig. S7F. H) Quantification of cCas3+ area from entire sections revealed increased apoptotic areas in +Dox (n=6) compared to −Dox tumors (n=5), **p<0.01. I) Flow cytometry of proliferative (Ki67+) cells in syngeneic B16F1 tumors expressing caALK4 (+Dox) or not (−Dox). A trend for decreased proliferation in caALK4 tumors (n=6) compared to controls (n=4) was not significant (p=0.09).

### Activin-A-dependent B16F1 melanoma growth is mediated by impaired tumor immune surveillance

To test whether paracrine Activin-A signaling promotes tumor progression by inhibiting anti-tumor immunity, B16F1 cells were inoculated subcutaneously or intravenously into FoxN1^nu/nu^ that lack functional T cells, or into Rag1^−/−^ mice devoid of V-D-J recombination in antigen receptors. In sharp contrast to immune-competent wild-type hosts, loss of either *FoxN1* or *Rag1* enabled CTRL tumors to grow as fast as those with sustained *INHβA* expression, even though Activin-A provoked cachexia irrespective of the host genotype (**Fig. 4A-F**). These results suggest that Activin-A accelerates B16F1 melanoma growth by blunting T cell-mediated anti-tumor immunity. In good agreement, the same lentiviral *INHβA* transgene also failed to stimulate tumor growth or experimental lung metastases in xenografts of human melanoma cell lines in immunodeficient FoxN1^nu/nu^ mice (**Fig. S5D-G**).

**Figure 4.**
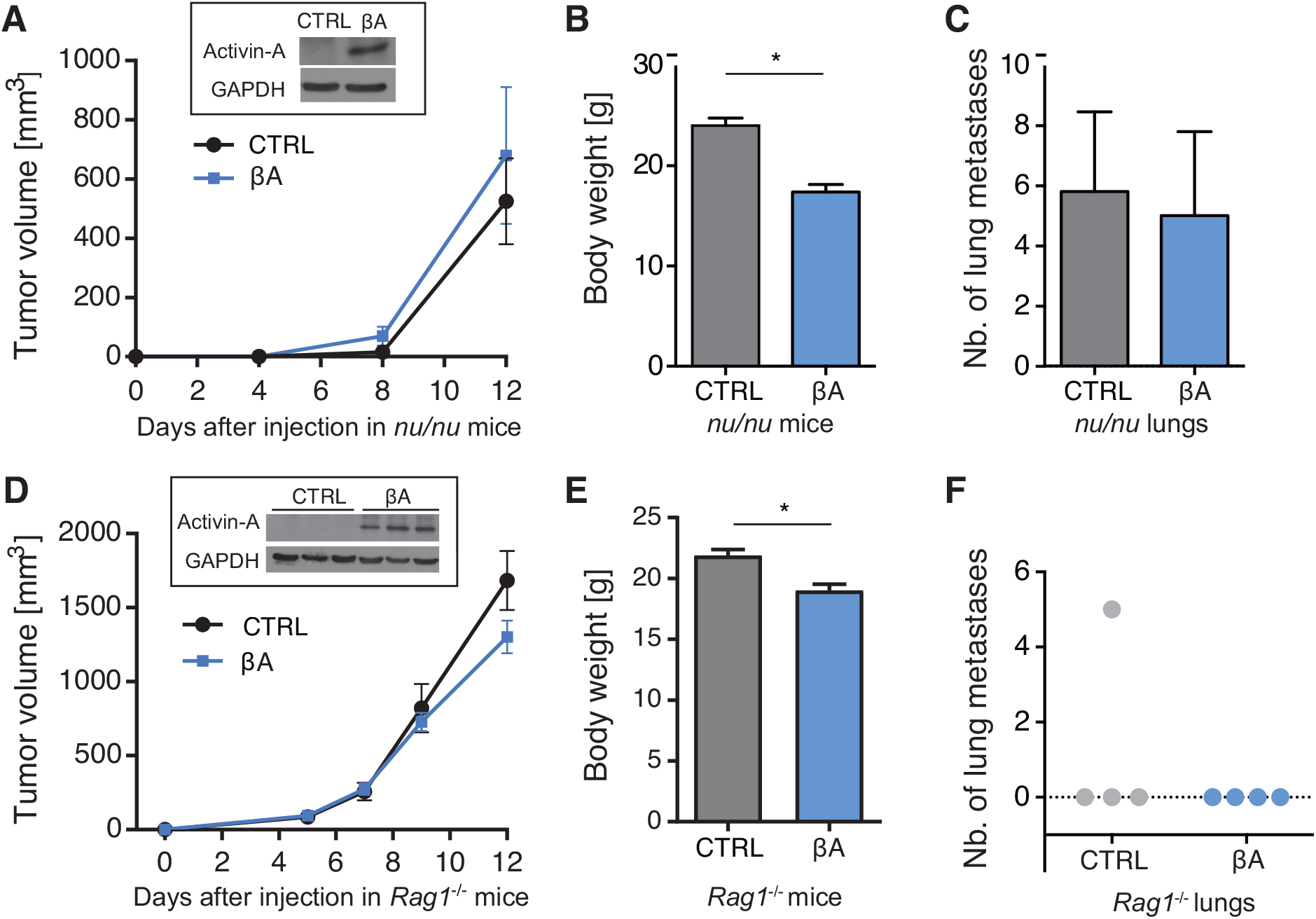
Tumorigenic activity of paracrine Activin-A signaling is mediated by the adaptive immune system. A) Growth curves of CTRL or βA-expressing B16F1:GFP tumors (n=5 each) after intradermal injection in the right flank of Foxn1^nu/nu^ mice. Inset: Western blot of Activin-A in randomly chosen tumor extracts at the experimental endpoint. GAPDH served as loading control. B) Body weights of the hosts of the tumors shown in (A) (mean±SEM, *p<0.05). C) Experimental lung metastases observed after tail vein injection of CTRL or βA-expressing B16F1:GFP (n=10 each) in Foxn1^nu/nu^ mice. D) Volumes of intradermal CTRL-(n=4) or βA-B16F1:GFP tumor grafts (n=3) in the right flank of *Rag1*^−/−^ mice. Western blots were as in panel A. E) Body weight at the time of tumor resection. Rag1^−/−^ mice injected with βA expressing B16F1:GFP develop cachexia (n=4) compared to CTRL injected animals (n=4). Data represent mean±SEM; ***p<0.001 F) Numbers of experimental lung metastases after tail vein injection of B16F1:GFP ± CTRL (n=4) or βA (n=4) in *Rag1*^−/−^ mice.

### Angiogenesis is enhanced by Activin-A but is not sufficient to promote B16F1 melanoma growth

Depending on the context, Activin-A signaling may also promote or inhibit angiogenesis which can be rate-limiting for tumor oxygenation and nutrient supply (Lewis et al., 2016). To assess whether increased tumor growth correlates with increased tumor vascularization, we labelled blood vessels in thick cryosections of syngeneic B16F1 grafts using CD31 antibodies. Quantification in z-stack reconstructions of entire sections of 15 CTRL and 14 INHβA tumors showed that Activin-A significantly increased the vascular density (**Fig. 5A**). Conversely, pimonidazole staining of hypoxic areas in the same sections was 4-fold reduced, indicating that Activin-induced blood vessels were functional (**Fig. S8A, B**). Since Activin-A has been shown to inhibit endothelial cell growth and tubule formation *in vitro* (Kaneda et al., 2011), we asked whether its pro-angiogenic effect in B16F1 melanoma could be mediated by macrophages. However, FACS sorting of dissociated B16F1 melanoma at the experimental endpoint revealed no changes in total myeloid populations, and M2-like macrophages marked by CD206 staining were decreased in *INHβA* compared to CTRL tumors (**Fig. S8C**). Also in human melanoma with the highest levels of Activin-A, markers of lymphocytic or myeloid infiltrates appeared to be reduced rather than increased (**Fig. S9**), although recruitment of such infiltrates or their functions may vary depending on Activin-A signaling strength or duration. Interestingly, angiogenesis was similarly increased in Rag1^−/−^ hosts were tumor growth remained unchanged upon *INHβA* overexpression, suggesting that increased vascularization alone cannot account for the tumorigenic effects of Activin-A (**Fig. 5B**).

**Figure 5.**
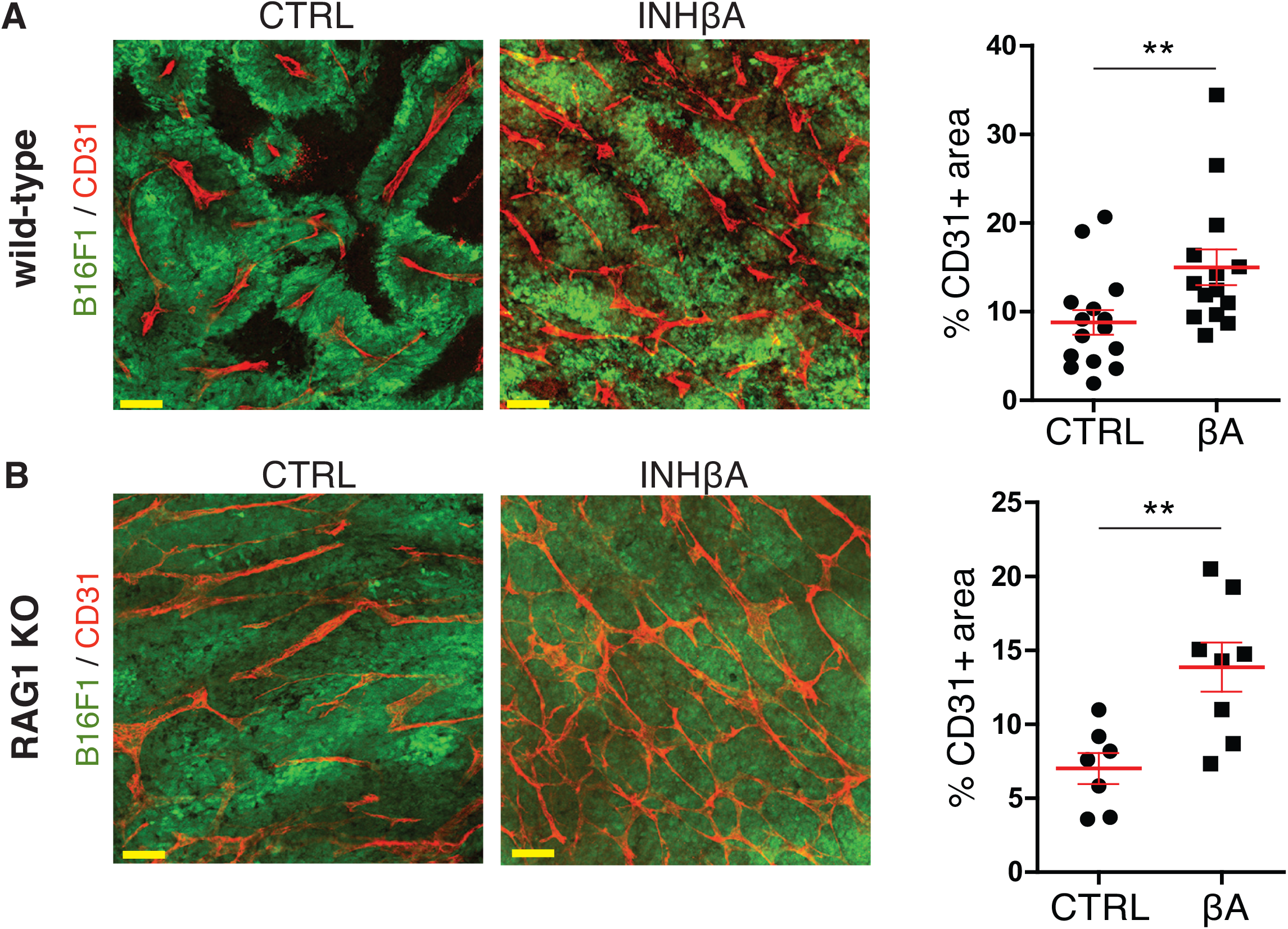
Paracrine Activin-A can stimulate tumor angiogenesis and oxygenation independently of adaptive immuity. CD31 immunofluorescence staining (red) of thick sections of B16F1:GFP CTRL or βA syngeneic intradermal grafts (green) in wild-type and Rag1^−/−^ C57BL/6 mice. Quantifications (right panels) show that βA-expressing tumors had significantly increased vascular coverage compared to CTRL tumors both in wild-type (βA: n=14; CTRL: n=15) and in Rag1^−/−^ hosts (βA: n=8; CTRL: n=7).

## DISCUSSION

Previous studies reported both tumor-suppressive and oncogenic effects of Activin-A, but a role in anti-tumor immunity and the relative contributions of autocrine versus paracrine signaling in an *in vivo* model of melanoma remained to be evaluated. Here, we show that the net outcome of a gain in paracrine Activin-A signaling in mouse and human melanoma grafts is determined by whether or not the tumor host has functional adaptive immunity. Tumorigenic paracrine effects on adaptive immunity trumped pro-apoptotic autocrine signals within cancer cells to overall facilitate primary and metastatic growth. Ectopic Activin-A signaling also stimulated tumor vascularization and, concurring with previous reports, systemic cachexia, and these effects were preserved in *Rag1*^−/−^ mice albeit without accelerating tumor growth. Thus, a potential boost in nutrient supply by recycled tissue breakdown products was either insufficient to fuel tumorigenesis or neutralized by growth-inhibitory autocrine Activin receptor signaling within tumor cells. To our knowledge, these results furnish the first direct evidence that adaptive immunity is required for a tumorigenic Activin function, and that autocrine and paracrine signaling mediate opposite effects on melanoma growth *in vivo.* Based on these findings, future strategies to boost the efficacy of immunotherapies should consider targeting Activin-A.

### Activin-A only stimulates melanoma growth in mice that have functional T cells

Our main finding is that paracrine Activin-A signaling was tumorigenic specifically in immunocompetent hosts, despite a proapoptotic function of autocrine Activin/Nodal receptor signaling within melanoma cells. Thus, a lentiviral *INHβA* transgene encoding secreted Activin-A in syngeneic B16F1 mouse melanoma grafts increased intradermal tumor growth and the frequency of experimental lung metastases specifically in wild-type C57BL/6 mice. By contrast, in syngeneic *Rag1*^−/−^ hosts that lack T- and B-lymphocytes, or in athymic *FoxN1*^nu/nu^ mice devoid of only T cells, neither blockade of endogenous Activin/Nodal receptor ligands in the human C8161 melanoma cell line nor overexpression of Activin-A revealed a tumorigenic function. Technical artifacts arising from clonal variation or specific cell lines are unlikely because we used non-clonal pools of lentivirally transduced cells, and reproducible results were obtained with multiple independent batches of cells and with additional human melanoma cell lines. In keeping with our observations, also other tumor types in immunocompromised mice revealed minimal or no oncogenic effects of Activin-A (Zhou et al., 2010). Notable exceptions are pancreatic and esophageal cancer and R30C mammary carcinoma cells where gain of autocrine Activin signaling directly stimulates tumor cell stemness or survival (Krneta et al., 2006, Lonardo et al., 2011).

Induction of metastatic disease in chemically induced squamous cell carcinoma by a keratinocyte-specific *INHβA* transgene facilitates infiltration by immunosuppressive Tregs while reducing the number of resident γδ-TCR-positive dendritic epidermal T lymphocytes (DETC), consistent with a potential role in promoting immune evasion (Antsiferova et al., 2011). Depletion of CD4-positive T cells, including Tregs did not suppress the tumorigenicity of transgenic Activin-A in this skin carcinoma model (Antsiferova et al., 2016). However, it will be interesting to compare in future studies the potential of Tregs or tumor-infiltrating cytotoxic T-lymphocytes or other T cell subsets to mediate effects of Activin-A on anti-tumor immunity in melanoma. A role in suppressing anti-tumor immunity may explain why *INHβA* expression is more frequently upregulated in human melanoma and other solid human tumors than expected for a neutral bystander (Hoda et al., 2016, Wu et al., 2015).

### Inhibition of melanoma growth by autocrine signaling may curtail oncogenic effects of secreted Activin-A on the tumor microenvironment

Even though Activin receptor signaling inhibits cell proliferation and induces apoptosis in normal melanocytes (Stove et al., 2004), treatment with recombinant Activin-A did not inhibit the proliferation or viability of cultured B16F1 cells *in vitro.* To more directly assess a potential role for autocrine Activin/Nodal signaling in the B16F1 model, we introduced a doxycycline-inducible ligand-independent mutant ALK4 transgene. We found that autocrine ALK4 signaling inhibited B16F1 cell proliferation and tumor growth rather than stimulating it. This confirms that oncogenic Activin-A activity in immunocompetent syngeneic hosts was mediated by paracrine signaling. Since melanoma cell survival was impaired by caALK4 but not by secreted Activin-A, autocrine signaling activity of the ligand is likely attenuated. In the hepatocyte lineage, autocrine anti-proliferative Activin-A signaling is frequently attenuated by secreted antagonists in hepatocarcinoma (reviewed in Deli et al., 2008). Also in human melanoma, dynamic changes in the levels of FST expression during progression of melanoma *in situ* to metastatic growth may modulate Activin-A responses (Heinz et al., 2015, Stove et al., 2004). However, whether FST influences the balance between autocrine and paracrine Activin signaling remains to be determined.

### Interplay of tumor growth and cachexia

Commonly associated with advanced cancer in human patients, cachexia reduces life quality and drug responses while increasing morbidity and mortality (Fearon et al., 2013). In pancreatic MIA PaCa-2 xenografts, a comparison of tumor growth with the time course of Activin-induced cachexia suggests that associated metabolic changes or general weakening curtails a growth-promoting effect of autocrine Activin signaling (Togashi et al., 2015). If cachexia similarly curbs the growth of B16F1 melanoma growth, tumor growth should slow down concurring with the onset of Activin-induced cachexia at least in immunodeficient hosts. Such a trend in *Rag1*^−/−^ hosts was not statistically significant and not seen in *nu/nu* hosts. The observed tumor growth curves also do not support a model that cachexia is rate-limiting for the supply of essential amino acids and other metabolites to cancer cells in a process of ‘auto-cannibalism’ (Theologides, 1979).

### Activin-induced tumor angiogenesis and its uncoupling from tumor growth

We found that paracrine Activin-A signaling in the syngeneic B16F1 melanoma model also stimulated tumor vascularization. However, a similar increase of blood vessels in *Rag1*^−/−^ hosts was not sufficient to facilitate tumor growth. Although we could not stain enough tumors in *Rag1*^−/−^ mice with pimonidazole to formally exclude a stimulatory effect of adaptive immunity on vessel functionality and hypoxia, it is interesting to note that tumor growth was also uncoupled from angiogenesis in immunodeficient SCID mice bearing mammary carcinoma xenografts, where gain of Activin-A signaling diminished tumor angiogenesis without affecting vascular perfusion (Krneta et al., 2006). Interestingly, however, Activin signaling within endothelial cells in this breast cancer model and other tumors was cytostatic and reduced sprouting and blood vessel density (Breit et al., 2000, Kaneda et al., 2011, Krneta et al., 2006), indicating that proangiogenic activity is likely indirect. T cell-derived cytokines are unlikely involved since Activin-A stimulated the vascularization of B16F1 tumors even in *Rag1*^−/−^ syngeneic hosts. Possibly, class I inflammatory macrophages that stimulate vascularization in Activin-induced skin squamous cell carcinoma mediate proangiogenic activity (Antsiferova et al., 2016). Although, the total number of infiltrating macrophages was unchanged by Activin-A in B16F1 melanoma under the conditions examined, and since increased angiogenesis was not sufficient to promote tumor growth, we did not further investigate whether Activin-A directly polarized a proangiogenic macrophage subtype in this model.

Overall, our findings suggest that paracrine Activin-A should be considered as a novel target for personalized therapies not only to reduce cachexia and melanoma vascularization, but also to enhance the efficacy of immunotherapies.

## MATERIALS AND METHODS

### Melanoma grafts

1×10^6^ B16F1 cells were injected intradermally into the right flank of 8-12 week old female wild-type (Harlan) or Rag1^−/−^ C57BL/6 (EPFL animal core facility) syngeneic hosts, or of Hsd-athymic *nu/nu* mice (Harlan). Animal body weights and tumor sizes were measured every two days. Tumor volumes were calculated using the formula *length x width x depth.* Experimental lung metastases were obtained by injecting 3×10^5^ B16F1 cells into the tail vein. Pigmented metastases visible at the surface of each lung lobe were counted 3 weeks after injection. Where indicated, animals were fed with chow containing 0.625 g/kg doxycycline (Provimi Kliba AG, Switzerland). All procedures were according with Swiss legislation and approved by the cantonal veterinary adminstration. Generation of cell lines and *in vitro* assays are further documented in Supplemental Methods.

### Gene expression analysis

Total RNA from melanoma cell lines and tumors was isolated using Trizol reagent (Sigma or ThermoFisher) and guanidinium/CsCl gradient, respectively, or by using RNeasyMini kit (Qiagen). 1 μg of total RNA was used for cDNA synthesis using the Superscript III Reverse Transcription Kit (Invitrogen). Quantitative polymerase chain reaction (qPCR) assays were performed using SYBR^®^ green chemistry according to manufacturers’ instructions (Applied Bioscience), or commercial Taqman probe for INHβA (Eurogentec). PCR primer sequences are listed in Table S1.

### Cell cycle and Ki67 analysis

After treatment with 250 μg/ml doxycycline for 24, 48 or 72 hours in 6-well plates, cells were incubated with 10 μM EdU for 30 min and then trypsinized, washed with PBS, and fixed for 20 min with 4% PFA, permeabilized with 0.5% Triton X-100 in PBS and stained for 30 minutes with Alexa-647-coupled azide in presence of copper sulfate and sodium ascorbate. Cells were then washed and stained with DAPI and analyzed using a CyAn ADP analyzer (Beckman Coulter). For Ki67 immunostaining, tumors were dissociated into single cells in 0.02 mg/ml DNase I and 1 mg/ml collagenase mix (Sigma) using a GentleMACS tissue octo dissociator (Miltenyi). Dissociated cells were stained with live and dead blue dye (Life Technologies), washed and labelled with antibodies against CD45-APC-Vio770 (Miltenyi, 130-105-463), CD31-BV605 (BD Biosciences, 740356), and CD140a-PE (eBioscience, 12-1401-81) to exclude leukocytes, endothelial cells and fibroblasts, respectively. After washing, cells were fixed and permeabilized using FoxP3 fix and perm buffer set (eBiosciences) and stained with eFluor 450-conjugated Ki67 antibodies (eBiosciences, 48-5698-80) using permeabilization buffer (eBiosciences). Samples were acquired using an LSRII cytometer (Becton Dickinson).

### Statistical analysis

All statistical analyses were performed using GraphPad Prism v6 software. Data were analyzed using Mann Whitney (for non-parametric data), T-tests, 1-way or 2-way ANOVA with Bonferroni correction for parametric data. A p-value <0.05 was considered significant.

## ACKNOWLEDGEMENTS

We thank Dr. Joerg Huelsken for the kind gift of lentiviral Gateway cloning vectors, Dr. Malcom Whitman for HA-tagged human ALK4 cDNA, and Drs. U. Koch, J. Faget and M. De Palma for advice on FACS, and M. Langegger for culturing melanoma cells for zebrafish transplantations, and the EPFL School of Life Sciences technology cores and animal care facility. This work was supported by grants to D.B.C. from the Swiss Cancer League (KLS – 02487-08-2009 and the Swiss National Science foundation (31003A_156452).

## SUPPLEMENTAL DATA

Commercial antibodies against mature NODAL have been reported to stain metastatic but not primary melanoma, and a FACS-sorted subpopulation of metastatic C8161 melanoma cells transfected with a NODAL antisense morpholino displayed reduced tumor growth compared to control cells when grafted into immunodeficient *nu/nu* mice (Postovit et al., 2008, Topczewska et al., 2006). However, a 35 kDa protein detected in extracts of C8161 melanoma cells (Hardy et al., 2010, Postovit et al., 2008, Topczewska et al., 2006) did not correspond in size to the known molecular weights of NODAL precursor (39-41 kDa) or processed form (12 kDa) (Constam and Robertson, 1999, Le Good et al., 2005). Among potential splice variants (Strizzi et al., 2012), only two transcripts share exons 2 and 3 encoding a functional NODAL precursor that can react with antibodies against the mature region (**Fig. S2A**). Therefore, to monitor *NODAL* mRNA expression, we designed intron-spanning primers in exons 2 and 3 which detect correctly spliced mRNAs in fetal brain or glioblastoma control samples. Unexpectedly, however, we did not detect spliced *NODAL* transcripts in C8161 cells or in any of the melanoma cell lines or 24 human melanoma biopsies analyzed, including C8161 cells, even after as many as 35 PCR cycles (**Figs. 1B, S2B**). Others have argued that correctly spliced *NODAL* mRNA can only be detected with special technical expertise using primer sequences that were not disclosed (Strizzi et al., 2012). However, a technical problem with our primers is unlikely since they amplified spliced *NODAL* mRNA fragment of the correct size (288 bp) both in fetal brain and glioblastoma total RNA (**Fig. S2B**).

To analyze NODAL protein, we raised a custom antibody against the fully conserved peptide KQYNAYRCEGECPNPV of human NODAL (**Fig. S2C**). In Western blots of transfected HEK293T and C8161 cells transduced with Nodal lentivirus (positive controls), the custom antiserum reacted with these recombinant control proteins as expected. However, no specific band was detected either in fresh lysates of nontransfected C8161 cells or in the less aggressive C81-61 cell line or their conditioned media (**Fig. S2D**, and data not shown). Species differences of human NODAL cannot account for the absence of specific bands since mature mouse Nodal is 98% identical and the antigenic peptide is 100% conserved (**Fig. S2C**). Also in the prodomain, NODAL is only 6 amino acids shorter in human than in mouse, and a single N-glycosylation motif (NWT) that is functional in mouse Nodal (Blanchet et al., 2008) is conserved.

To address the discrepancy between these results and published data, we compared our custom antibody with the commercial antibodies from Santa Cruz or R&D that were previously used to detect NODAL in melanoma (Hardy et al., 2010, Topczewska et al., 2006). While both of them recognized recombinant Nodal in HEK293T cells, they were less sensitive than our custom antiserum both on stripped (**Fig. S2D**) and on non-stripped blots (not shown). Furthermore, superposition of blots revealed that each commercial antibody also cross-reacted with distinct sets of additional bands around 35 kDa and above 49 kDa, especially if cells were extracted with detergent (Postovit et al., 2008) instead of Laemmli buffer (**Fig. S2E**). Importantly, however, we also observed these bands in MyoEP cell extracts (negative control, courtesy of Dr. Postovit), and proteins reacting with the Santa Cruz antibody did not overlap with those detected by R&D antibody (**Fig. S2E**). Therefore, and since no endogenous NODAL was detected with the more sensitive custom Nodal antibody, it seems that both commercial antibodies each cross-react with distinct non-specific proteins.

To distinguish whether C8161 cells secrete NODAL or a related activity, we tested the potential of conditioned medium to induce the SMAD3 luciferase reporter CAGA-Luc in HEK293T cell lines that express the Nodal co-receptor Cripto. As a negative control, we also used analogous reporter cells stably transduced with the related protein Cryptic. As shown previously (Fuerer et al., 2014), only Cripto but not Cryptic mediate induction of luciferase expression by Nodal, whereas treatment with recombinant Activin-A potently induced the reporter in both cell lines (**Fig. S2F**). Importantly, CAGA-Luc was also induced by C8161-conditioned medium independently of Cripto, and this activity was sensitive to inhibition by the Activin-specific antagonist Follistatin which does not inhibit NODAL (**S2F**). Concurring with the above RT-PCR and Western blot data, these results indicate that C8161 cells do not secrete detectable amounts of Nodal activity under the conditions examined. To directly validate whether C8161 secrete Activin-A, we also analyzed cell lysates and conditioned medium by Western blot. Using an antibody that reacts only with secreted Activin-A dimers, we detected mature Activin-A in its processed form specifically in conditioned medium of C8161 and Me300 cells, but not in the medium of C81-61 or Me290 melanoma cells or in cell lysates (**Fig. S2G**). Taken together with our zebrafish experiments (**Fig. S4**), these results show that C8161 cells under the conditions examined secrete biologically active Activin-A but not detectable amounts of NODAL.

## SUPPLEMENTAL MATERIALS AND METHODS

### Cell lines

B16F1 melanoma cells (ATCC) were maintained in Dulbecco’s modified Eagle’s medium supplemented with 10% fetal bovine serum and penicillin-streptomycin (GIBCO Invitrogen, Grand Island, NY). Melanoma cell lines SK-Mel-37, NA8-MEL, and MNT-1 were kind gifts from Drs. Y.T. Chen (Ludwig Institute for Cancer Research, New York Branch, New York, NY), F. Jotereau (U211, Institut National de la Santé et de laRecherche Médicale, Nantes, France), and P. G. Natali (Istituto Regina Elena, Rome,Italy), respectively. Metastatic human C8161 melanoma cells and non-invasive related C81-61 cell line were kindly provided by Dr. Mary Hendrix. C81-61 culture medium was Ham’s F10 (Invitrogen) supplemented with 15% fetal bovine serum, 1% gentamicin and MITO+ serum extender (BD Biosciences). C8161 cells were cultured in RPMI medium supplemented with 10% fetal bovine serum and 1% gentamicin (all from Invitrogen). The same medium was used for all other melanoma cell lines that were established from human cutaneous, visceral, or lymph node metastases, except for Me300, Me191 and T921A, which were from primary tumors. Human HepG2 and HEK293T cell lines were from ATCC (Middlesex, UK). HepG2 reporter cells stably transduced with lentiviral CAGA-Luc reporter and a Renilla luciferase for normalisation and with Cripto or Cryptic have been described (Fuerer et al., 2014). All cell lines were tested before *in vivo* use and found to be free of *Mycoplasma* (SouthernBiotech, 13100-01).

### Expression vectors

The lentiviral vector pRRL-EF1a-MCS-SV40puro was derived from pRRLSIN.cPPT.PGK-GFP.WPRE (Addgene ID 12252) by replacing PGK-GFP with the EF1a promoter of pEF1a-Myc-HIS (invitrogen) and with an MCS//SV40-Puro cassette. *INHβA* cDNA (nucleotides 218-1609 of the human reference sequence NM_002192.2) containing the myc epitope sequence GAGCAGAAGCTGATATCCGAGGAGGACCTG after position 1183 (kind gift of Tom Jessell, Columbia University, New York) was subcloned as an Spe I restriction fragment into the Xba I site of pRRL-EF1a-MCS-SV40puro. The extracellular domain of murine Acvr2b (nucleotides 369 to 770 of GenBank entry NM_007397.3) was fused via a PreScission cleavage site to the Fc fragment of human IgG1 (nucleotides 721-1401 of GenBank entry AB776838.1) and cloned as an Nhe I - Xba I fragment into the Xba I site of pRRL-EF1a-MCS-SV40puro. pLenti CMV rtTA3 Blast (w756-1) was a gift from Eric Campeau (Addgene 26429). To generate HA-tagged caALK4 lentivector, HA-tagged caALK4 cDNA fragment was flanked by attB1 and attB2 sites using the PCR primers fw: 5’GGGGACAAGTTTGTACAAAAAAGCAGGCTccaccATGGGAAGTAGTAAAAGTAAGCCA3’ and rv: 5’ GTTCCTGATTATGCTAGCCTCtaaACCCAGCTTTCTTGTACAAAGTGGTCCCC3’ and inserted into pDONR221 (gift from Dr. J. Huelsken, EPFL) using Gateway BP clonase II (Invitrogen 11789020), followed by transfer into cPPT-tetO-mCMV-attR-IRESeGFP-WPRE (gift from J. Huelsken, EPFL) using Gateway LR clonase II (Invitrogen, 11791020).

### Lentiviral transduction

To produce lentiviral particles, packaging and envelope protein plasmids CMVΔR8.74 (Addgene 22036) and pMD2.VSVg (Addgene 12259) were co-transfected with Nodal, mycActivinA (INHβA) or AIIB-Fc or empty pRRL-EF1a-MCS-SV40puro vector control (CTRL) in HEK293T cells as described at a ratio of 13:7:21 (Fuerer et al., 2014). Cell supernatants were sterile-filtered (0.45-μm) and used either directly or after 50’000 × g ultracentrifugation and resuspension with 0.1% bovine serum albumin in PBS to transduce 1.2×10^6^ B16F1 or C8161 melanoma cells, followed by selection with puromycin (3 μg/ml). Doxycycline-inducible B16F1 cell lines were obtained by sequential transduction of lentiviral rtTA and HA-tagged caALK4, followed by selection with 5 μg/ml blasticidin and 1 μg/ml puromycin, respectively.

### Zebrafish assays

Human MycActivinA (INHβA) and Nodal cDNA (IMAGE clone 8327493) were cloned into pCS2+. MycActivinA was cloned into EcoRI and XbaI sites; the coding sequence is flanked by ActivinA 5’ and 3’ UTR sequences. A Myc tag was inserted between the pro- and mature domains. An 1.31 kb EcoR1 cDNA fragment encoding Human Nodal (IMAGE clone 8327493) was cloned into pCS2+. The pCS2+ zebrafish Squint construct was previously described (Muller et al., 2012). Plasmids were linearized with NotI (NEB), purified using a Wizard SV Gel and PCR Clean-Up System (Promega), and capped mRNA was generated from linearized plasmid using an SP6 mMessage mMachine kit (Ambion) followed by purification (RNAeasy kit, Qiagen). 50 pg of mRNA in 1 nl of RNAse-free water were injected into *MZoep* or wild type (TE strain) zebrafish embryos at the one-cell stage, followed by incubation at 28°C. Triplicates of 5 embryos each per condition were collected at shield stage for RT-qPCR analysis of the Nodal target gene *gsc* (F: 5’-gagacgacaccgaaccattt-3’, R: 5’-cctctgacgacgaccttttc-3’) and the house keeping gene *eF1alpha* (F: 5’-agaaggaagccgctgagatgg-3’, R: 5’-tccgttcttggagataccagcc-3’) as described (Muller et al., 2012). Total RNA was purified using NucleoZol extraction (Machery-Nagel), cDNA generated using a Superscript III Supermix kit, and RT-qPCR performed using a Platinum SYBR Green qPCR SuperMix-UDG kit (Invitrogen) on a BioRad CFX-Connect cycler.

For cell transplantations, 50-100 melanoma cells (≈500 pl of a cell suspension with 10^8^ cells/ml in Ca^2+^-free Hank’s medium) were grafted into pronase-dechorionated wild-type (TE) or *MZoep* embryos at sphere stage as described (Topczewska et al., 2006). After incubation at 31°C for 3 hours until shield stage, embryos were fixed and stained for *ntl* and *gsc* by whole mount *in situ* hybridization (Muller et al., 2012). Embryos were imaged using an Axio Zoom.V16 microscope (ZEISS).

### Reporter assays

HepG2 CAGA-Luc reporter cells stably expressing the Nodal co-receptor Cripto as well as Renilla luciferase for signal normalisation (Fuerer et al., 2014) were plated in 96-well plates (10^4^ cells per well). To monitor the activity of secreted Activin-A, conditioned media of melanoma cell lines were preincubated with or without 250 ng/ml Follistatin (R&D Systems) prior to addition to HepG2 CAGA-Luc reporter cells. Alternatively, as a positive control, reporter cells were directly transfected with Activin-A (INHβA) or empty control (CTRL) plasmids prior to plating, or treated with the indicated concentrations of recombinant Activin-A or Nodal (R&D Systems). After 24 hours, cells were extracted for 20 min in 100 μl potassium phosphate buffer containing 0.5% Triton X-100, and luciferase activities

### Scratch wound assay

Cells were plated in 6-well plates at confluent density of 4×10^6^ per well. After letting cells adhere for six hours, a scratch wound was made in the cell monolayer using a P200 pipette tip, and the place of the scratch was marked. The cells were washed with PBS and the medium was changed for the indicated treatments. Scratches were imaged at 0 hours and after 20 hours.

### Cell viability assay and melanin secretion

Where indicated, cells were preincubated with 250 μg/ml doxycycline for 12-24 hours. 5x10^3^ cells were then seeded in 96-well plates and cultured for 3 days in presence of doxycycline. Alamar blue reagent (Invitrogen, DAL1025) was added to wells 4 hours before fluorescence measurement at excitation and emission wave lengths of 560 and 590 nm, respectively. Melanin concentrations in conditioned medium from CTRL or INHβA expressing B16F1 cells were determined by measuring absorbance at 575 nm using a TECAN spectrophotometer.

### Whole mount staining

Tumors were fixed with 4% PFA overnight and cryoprotected with 20-30% sucrose in PBS and embedded in OCT. Free-floating thick cryosections (200 μm) were digested with proteinase K (20 μg/ml) in 10 mM Tris HCl buffer (pH 7.5) for 5 minutes at room temperature, followed by permeabilization in 100% MetOH for 30 minutes. Sections were washed three times with PBS and blocked with 3% skim milk in 0.3% Triton X-100, followed by overnight incubation with rat anti-CD31 monoclonal antibody (1:300, BD Pharmingen #553370). After rinsing three times with 0.3% Triton X-100 in PBS, sections were incubated with Alexa647-labelled secondary antibodies (1:800, Invitrogen A-21247). To monitor hypoxia, mice were injected intraperitoneally with pimonidazole (60 mg/kg of body weight; Hypoxyprobe, HP-7) 45 minutes before sacrifice with a lethal dose of pentobarbital (250 mg/kg). Hypoxic areas were stained with HP-Red549 antibodies (1:200, Hypoxyprobe, HP-7).

### Western blot analysis

Proteins in conditioned media were precipitated by adding 4 volumes of ice-cold acetone and incubation at -20°C for 12 hours. Washed pellets were resuspended in Laemmli buffer. Cell extracts were obtained by incubating cells in 20 mM Tris HCl, pH 7.5 containing 150 mM NaCl, 1 mM EDTA, 1 mM EGTA, 1% Triton, 2.5 mM sodium pyrophosphate, 1 mM β-glycerophosphate, 1 mM Na3VO4, 1 μg/ml leupeptin,complete protease inhibitor cocktail (Roche Diagnostics, Mannheim, Germany) and phosphatase inhibitor cocktail (Sigma, P5726) when assessing phosphoproteins. Protein concentrations were determined using Bradford assays (Bio-Rad, Munich, Germany). Proteins (20 μg/lane) in cell extracts were separated by 10% SDS–PAGE and transferred onto nitrocellulose membranes (Schleicher and Schüll, Dassel, Germany). After blocking for 1 h at room temperature with skim milk milk powder (5%) in phosphate-buffered saline and 0.1% Tween-20 (PBST), membranes were incubated overnight in PBST supplemented with 2.5% bovine serum albumin and primary antibodies against disulfide-linked Activin-A homodimers (#AF388 R&D, Düsseldorf, Germany) pSMAD2 (Cell Signaling 3108,), pSMAD2,3 (Cell Signaling 8828), HA-Tag (Sigma H-6908) or γ-tubulin (Sigma GTU88), each at 1:1000 dilution. Anti-NODAL antiserum (CAN3) was raised in rabbits against the peptide KQYNAYRCEGECPNPV (Eurogentec). After washing three times with PBS, secondary antibodies coupled to horseradish peroxidase were incubated at 1:5000 dilution and visualized after extensive rinsing with PBST using ECL chemiluminescence (Amersham Pharmacia).

### Detection of tumor associated macrophages

Tumors were dissociated in 4 ml Ca/Mg-free PBS containing 40 μl of Liberase DL (Roche, 28U/ml), 80 μl of Liberase TL (Roche, 14 U/ml), and 40 μl of DNase I (15 mg/ml, Sigma) for 45 min at 37°C. After filtration and red blood cell lysis, non-specific Fc binding was blocked using CD16/CD32 antibody (clone 93). Rinsed single cells (2×10^6^) were analysed on a BD LSRII cytometer after antibody and DAPI staining: Conjugated antibodies were anti-mouse CD45-ACP (eBioscience, 17-0451, 1:400), CD11b-PEcy-7 (eBioscience, 25-0112, 1:400), F4/80-APCcy7 (eBioscience, 14-4801, 1:400), CD11c-PE (eBioscience, 12-0114, 1:100), and CD206-biotin (BioLegend 14713, 1:50) combined with Streptavidin-PerCp/Cy55 (BioLegend, 405214, 1:300). Single stain and fluorescence minus one controls were used to determine specificity of staining and to set gates. TAMS were gated on viable CD45+ and F480+. M1-polarised macrophages were defined as CD45+F480+CD11c+CD206− whereas M2 were CD206+. Myeloid cells were defined as CD45+F480−.

## SUPPLEMENTAL FIGURE LEGENDS

**Figure S1.**
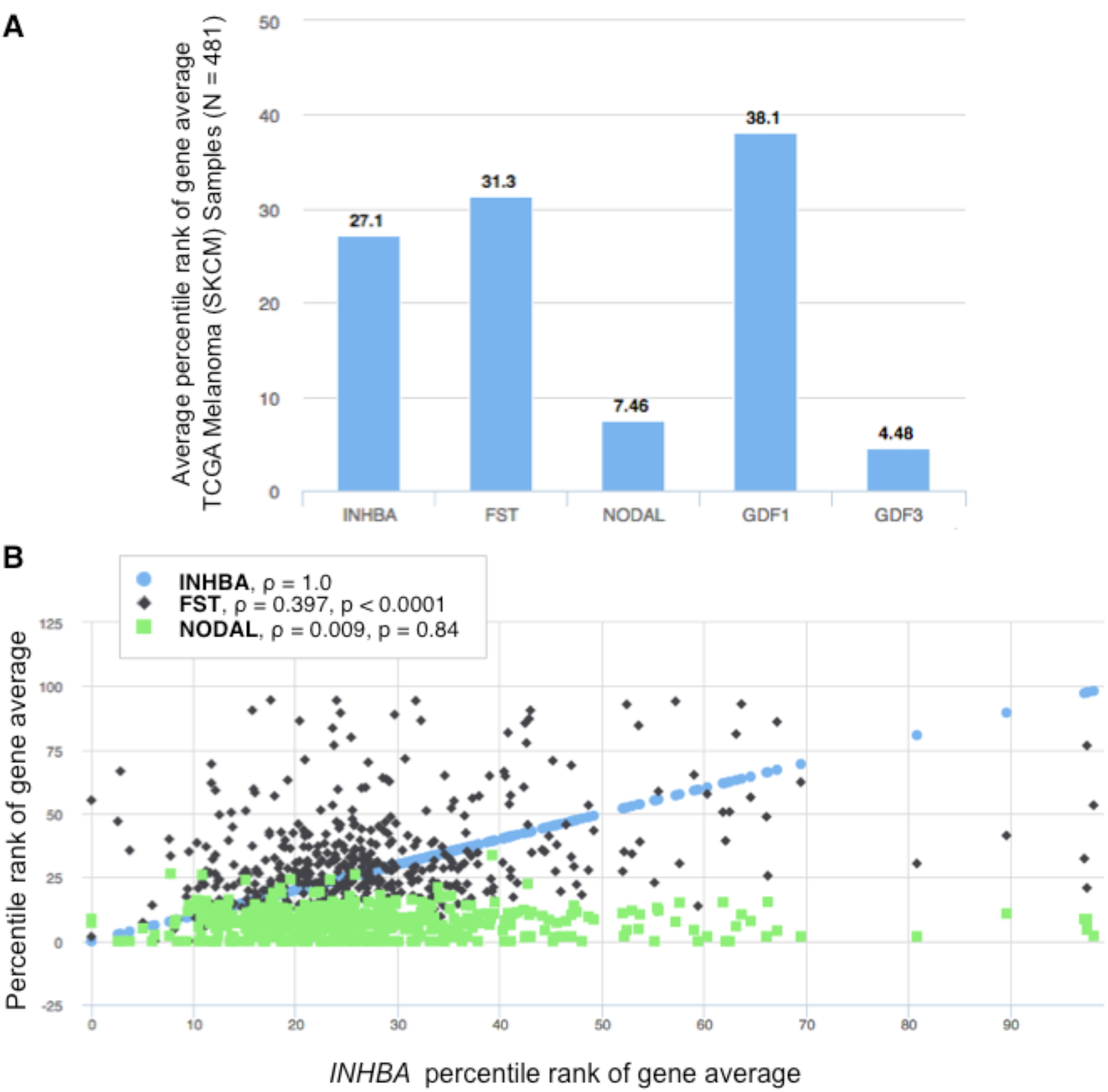
Expression of Activin receptor ligands and Follistatin in the TCGA Skin Cutaneous Melanoma RNAseq dataset. A) Average expression (percentile rank) of the indicated genes visualized using the UCSC Xenabrowser server (http://xena.ucsc.edu/getting-started/). B) Comparison of relative gene expression (percentile rank of gene average) in human melanoma plotted against the percentile rank of gene average of *INHβA* expression levels (x-axis). Spearman correlation (ρ) with *INHβA* mRNA expression and p-values (two-tailed) are indicated. Note that none of the tumors analyzed expresses high levels of *NODAL* mRNA (n=474).

**Figure S2.**
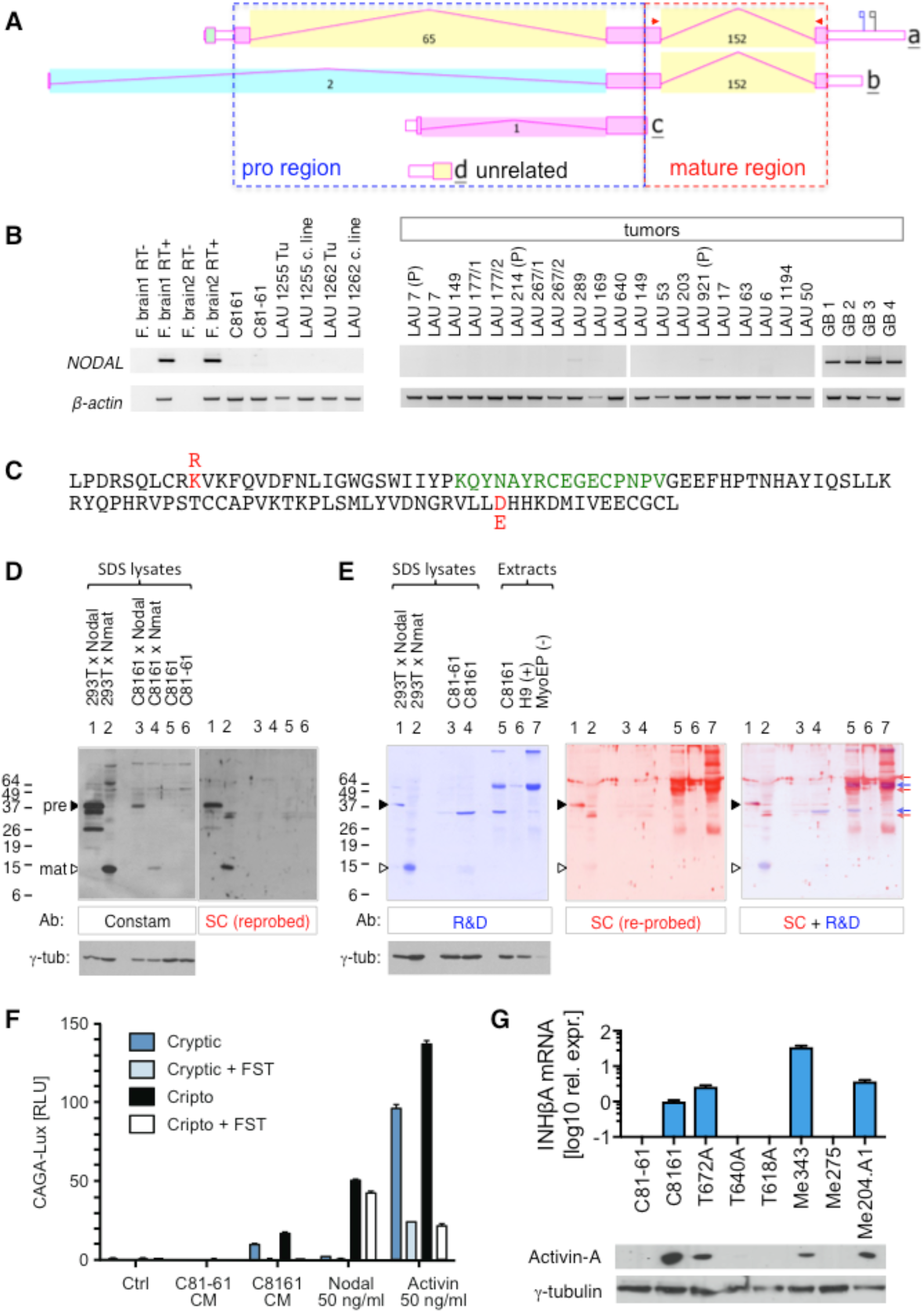
Analysis of NODAL mRNA and protein expression in human melanoma cell lines and tumors. A) NCBI AceView of human *NODAL* transcript variants *a* to *d*, representing 39 cDNA clones from diverse sources (*a*), 2 from brain and adult medulla (*b*), 1 from human ES cells (*c*), and one from a 3’-truncated incomplete cDNA of ES cells (**d**). Source: http://www.ncbi.nlm.nih.gov/IEB/Research/Acembly. Intron-spanning primers that detect all transcripts encoding functional protein (mature region) are indicated by red triangles. B) Reverse transcription PCR of NODAL mRNA in 22 human melanoma and in 4 glioblastoma (GB). Normal fetal brain (F.brain) cDNA samples are positive controls. C) Amino acid sequence of mature human NODAL protein. The homologous mouse sequence is identical except for two residues marked in red. The conserved immunogenic peptide used to raise custom NODAL antibody is highlighted (green). D) Western blot analysis of SDS lysates or detergent extracts prepared from the indicated cell lines (top) were probed with custom rabbit antiserum (Constam) before re-probing with commercial antibodies against mature NODAL from Santa Cruz (SC, H-110, right panel). Both antibodies react with full length Nodal (pre) and a constitutely mature mutant form (Nmat) in transfected HEK293T cells (Le Good et al., 2005). Custom Nodal antibody also detected the corresponding lentivirally transduced proteins in human C8161 cells. No NODAL protein was detected in non-transgenic parental C8161 cells or its non-metastatic counterpart C81-61. γ-tubulin served as loading control. E) Comparison of NODAL antibodies from R&D Systems AF1315 (blue) and Santa Cruz H-110 (red) by Western blot analysis of transfected HEK293T (control) and C8161 or C81-61 cell lysates. Superimposed image (right panel): Cross-reactivity of each antibody with several distinct proteins that do not co-migrate with uncleaved (lane 1) or mature Nodal (lane 2), except one faint band (open triangle) detected by R&D antibody that failed to react with the specific and more sensitive custom Nodal control antibody (panel D) or with Santa Cruz H-110. Reference extracts from C8161 cells (lane 5), human H9 embryonic stem cells (lane 6), or MyoEP cells (lane 7, negative control) were a courtesy of Dr. L. Postovit (Postovit et al., 2008). F) Induction of CAGA-Luc expression in stable HepG2 reproter cell lines by conditioned media of C81-61 (negative control) or C8161 cells treated with or without Follistatin (FST, 300 ng/ml), a specific antagonist of Activin-A that does not inhibit Nodal. Values represent mean±SEM of 2 independent experiments in triplicates. G) Taqman RT-qPCR of *INHβA* mRNA in C8161 cells and other human melanoma cell lines (top) and anti-Activin-A (*R&D* Systems, clone 69403) Western blot analysis of the corresponding conditioned media after non-reducing SDS-PAGE. The corresponding cell lysates probed with γ-tubulin antibody under reducing conditions are shown below. Data represent mean±SD of (n=2).

**Figure S3.**
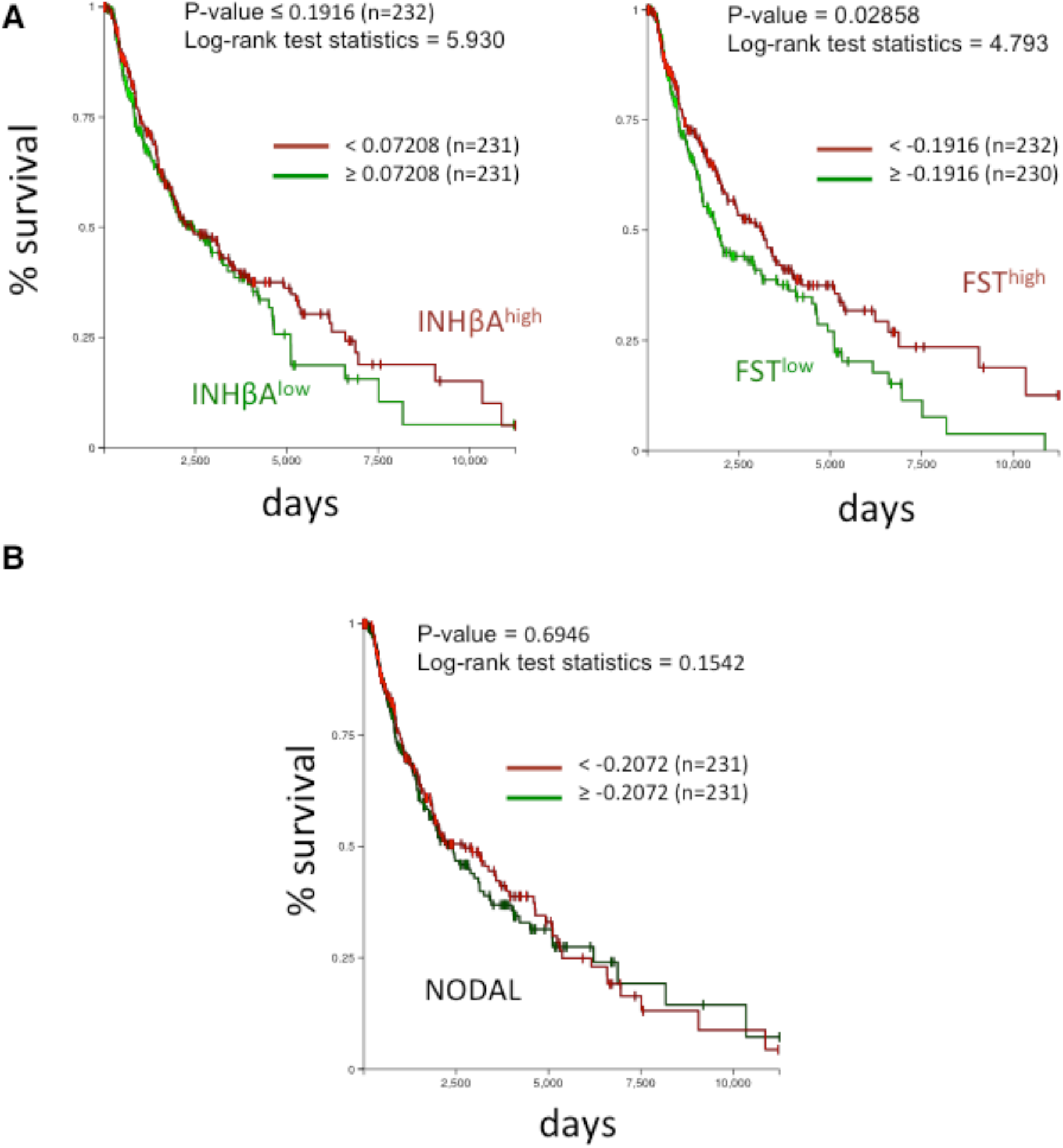
Survival of human patients in the TCGA Skin Cutaneous Melanoma dataset. A) Kaplan Meier plots reveal improved outcome for patients with high *Follistatin* expression (FST, p = 0.029), but not for low *INHBA* expression. B) Kaplan Meier plot of survival depending on relative levels of NODAL mRNA.

**Figure S4.**
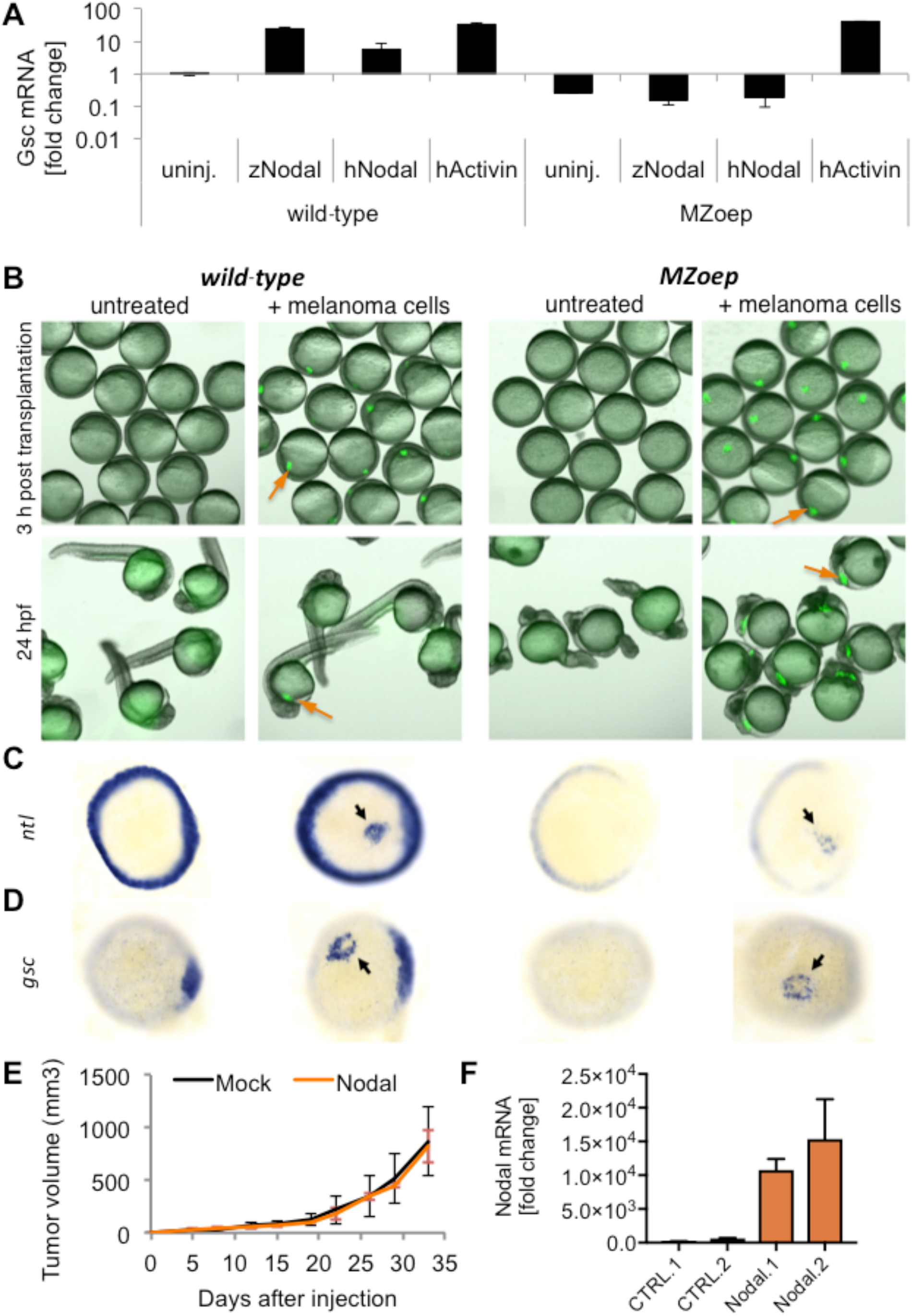
Nodal-independent induction of Smad2,3-mediated signaling and failure of Nodal to stimulate tumor growth by C8161 human melanoma cells. A) Wild-type and *MZoep* zebrafish embryos injected at the one-cell stage with 50 pg of mRNA encoding zebrafish Squint (zNodal), human NODAL (hNodal) or Myc-tagged Activin-A (Activin) were collected during early gastrulation for RT-qPCR analysis of the Nodal target gene *goosecoid* (*gsc*). While both human Nodal and Activin-A induced *gsc* in the wild-type, only Activin-A induced *gsc* in MZ*oep* embryos, demonstrating that human Nodal cannot activate Smad2,3 signaling in zebrafish independently of Oep, whereas Activin-A can. B) C8161-GFP human melanoma cells (top panel, arrows) were transplanted into wild-type and MZ*oep* sphere-stage zebrafish embryos. 24 h post fertilization (hpf), wild-type embryos exhibited generally normal development with some tissue protrusions, whereas MZ*oep* embryos exhibited the expected Nodal loss-of-function phenotypes. C, D) 3 h post transplantation, embryos were fixed for *in situ* hybridization of the Nodal target genes *no tail* (*ntl*, C) and *gsc* (D). Induction of both target genes around melanoma cell clones was evident in both wild-type and MZ*oep* embryos (arrows); additional blue staining indicates endogenous *ntl* expression around the circumference of embryos (C), or endogenous *gsc* expression on the dorsal side, respectively (D, right). Together with the RT-qPCR data, these results demonstrate Nodal-independent induction of Smad2,3-mediated signaling by C8161 human melanoma cells. Induction by C8161 grafts could be caused by Activin secretion (**Figs. 1C, D, S2F**). E) Growth curves of subcutanous human C8161 xenografts transduced with Nodal or control lentivirus (n=10 each). F) RT-qPCR analysis of Nodal mRNA in 2 randomly chosen tumors at the experimental endpoint (day 12). Error bars represent SD of triplicate PCR reactions.

**Figure S5.**
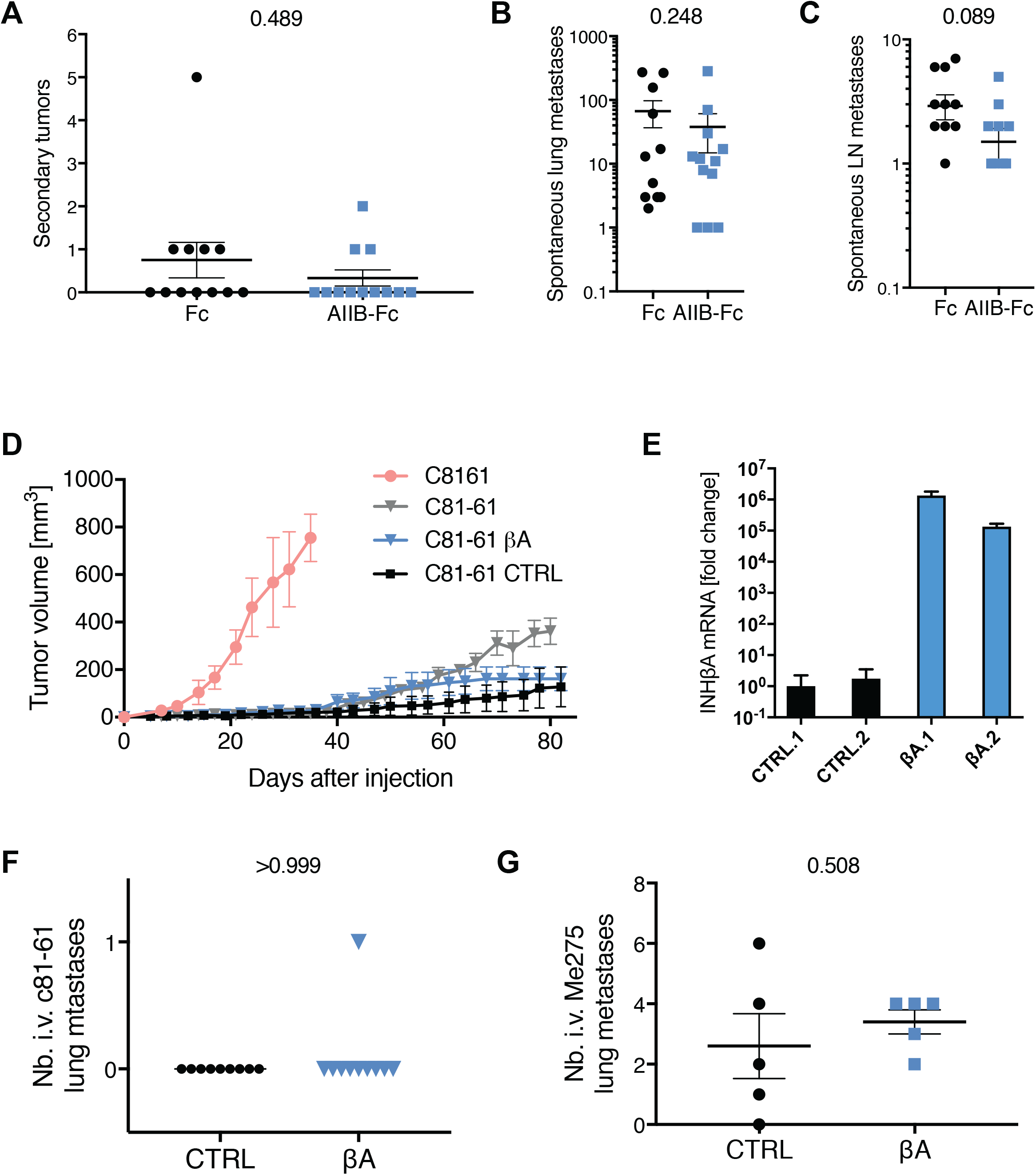
Analysis of human melanoma xenografts stably transduced with *AIIB-Fc* or *INHβA* transgenes. A-C) Lentiviral AIIB-Fc did not significantly inhibit spontaneous secondary C8161 melanoma growths 4 weeks after surgical removal of the primary s.c. graft (A, n=12), nor the numbers of spontaneous lung (B) or lymph node metastases (C) in FoxN1^nu/nu^ hosts. P-values are indicated at the top (two-tailed Mann Whitney). D) Stable transduction of *INHβA* lentivirus did not significantly accelerate the growth of C81-61 melanoma xenografts compared to CTRL (n=5 each; p=0.77 at endpoint). Growth curves of parental C81-61 cells and the related C8161 cell lines (n=5 each) expressing endogenous INHβA are shown for comparison. E) RT-qPCR analysis of relative *INHβA* expression levels in 2 randomly chosen C81-61 xenografts per group at the endpoint of the experiment. Data represent mean±SD of triplicates. F) Lentiviral *INHβA* expression did not induce lung metastasis in i.v. C81-61 grafts (βA: n=10, CTRL: n=9; p>0.99).

**Figure S6.**
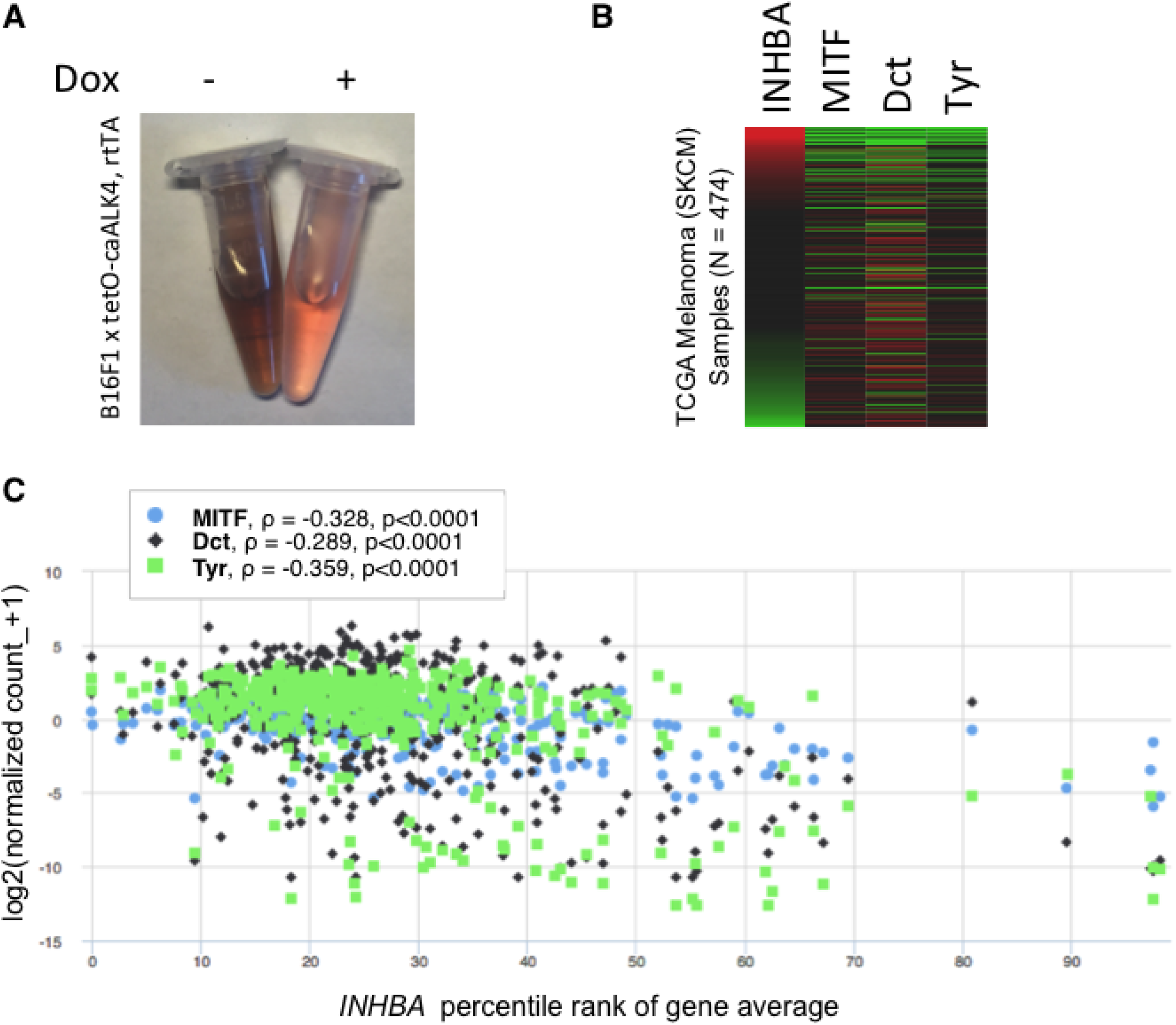
Correlation of INHBA status with markers of pigment synthesis in the TCGA Skin Cutaneous Melanoma dataset. A) Conditioned media of 1×10^5^ B16F1-caALK4 cells treated with or without doxycycline for 72 hours to induce caALK4 expression. Dark color indicates pigment secretion which was inhibited in caALK4 expressing cells. B) Heatmaps of *INHBA* expression and of the indicated markers of pigment synthesis in 474 human melanoma in the TCGA database analyzed using the Xenabrowser. C) Relative expression of the indicated pigmentation markers plotted against percentile rank of *INHβA* expression by Xenabrowser. Spearman correlations (ρ) with log2 values of normalized *INHβA* mRNA expression are indicated.

**Figure S7.**
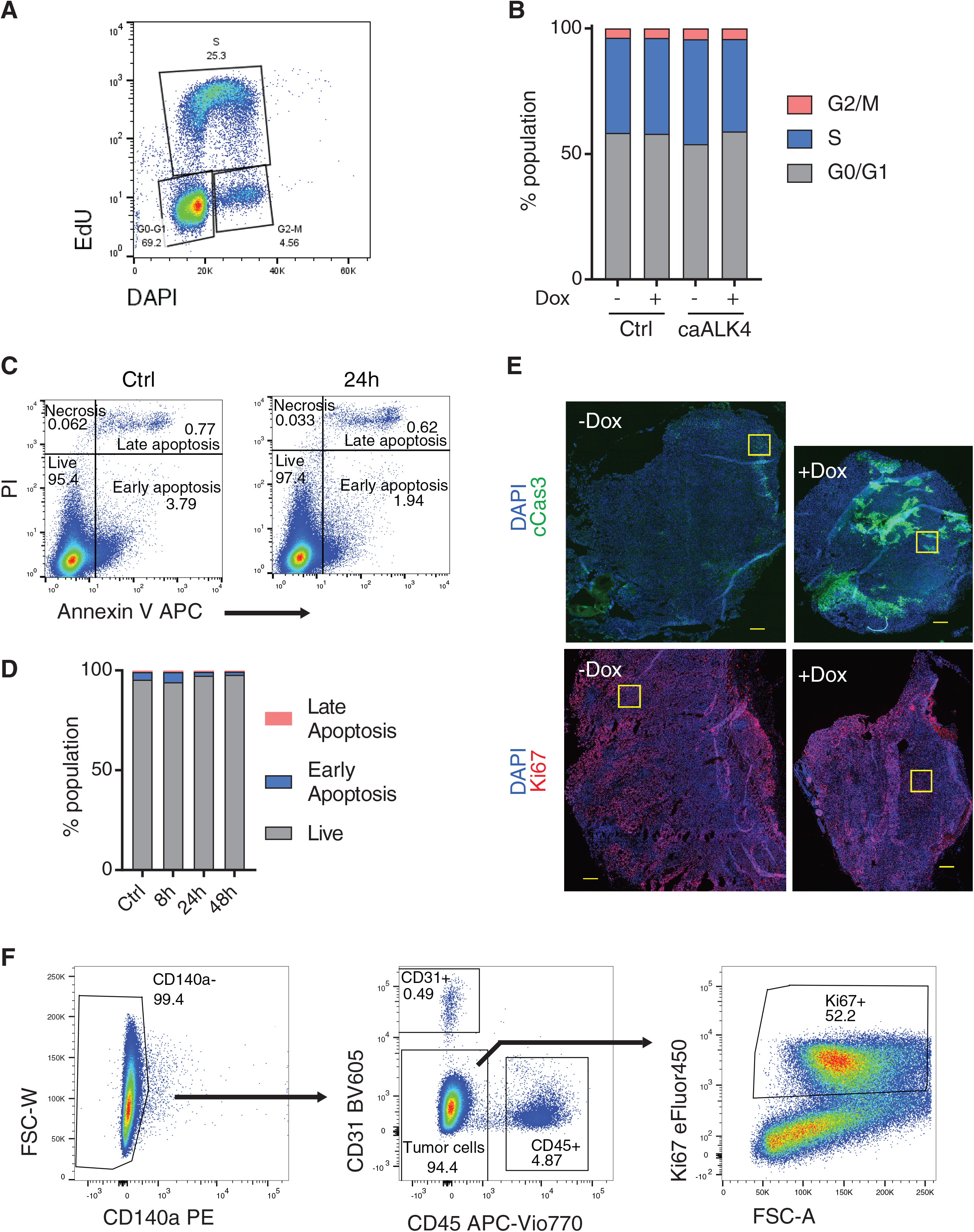
Analysis of B16F1 melanoma cell proliferation. A) A representative gating and B) flow cytometric quantification of DNA content (DAPI) and EdU-labelling of cultured B16F1 melanoma cells in G2/M- and S-phase after treatment with or without doxycycline for 48 hours. C) Representative gating of AnnexinV/propidium iodide (PI) stainings. D) Quantification of AnnexinV/PI assays after doxycycline treatment of inducible B16F1-caALK4 cells for 0, 8, 24, or 48 hours. E) Cleaved Caspase3 (cCas3) and Ki67 whole section immunofluorescence staining of syngeneic B16F1 tumors expressing caALK4 (+Dox) or not (-Dox). Corresponding magnifications are shown in **Fig. 3G**. Yellow squares represent the magnified areas. Scale bars = 250μm. F) Flow cytometric identification of Ki67-stained tumor cells (third panel) in B16F1 melanoma grafts after exclusion of CD140a+ (first panel), and CD31+ and CD45+ host cells (second panel).

**Figure S8.**
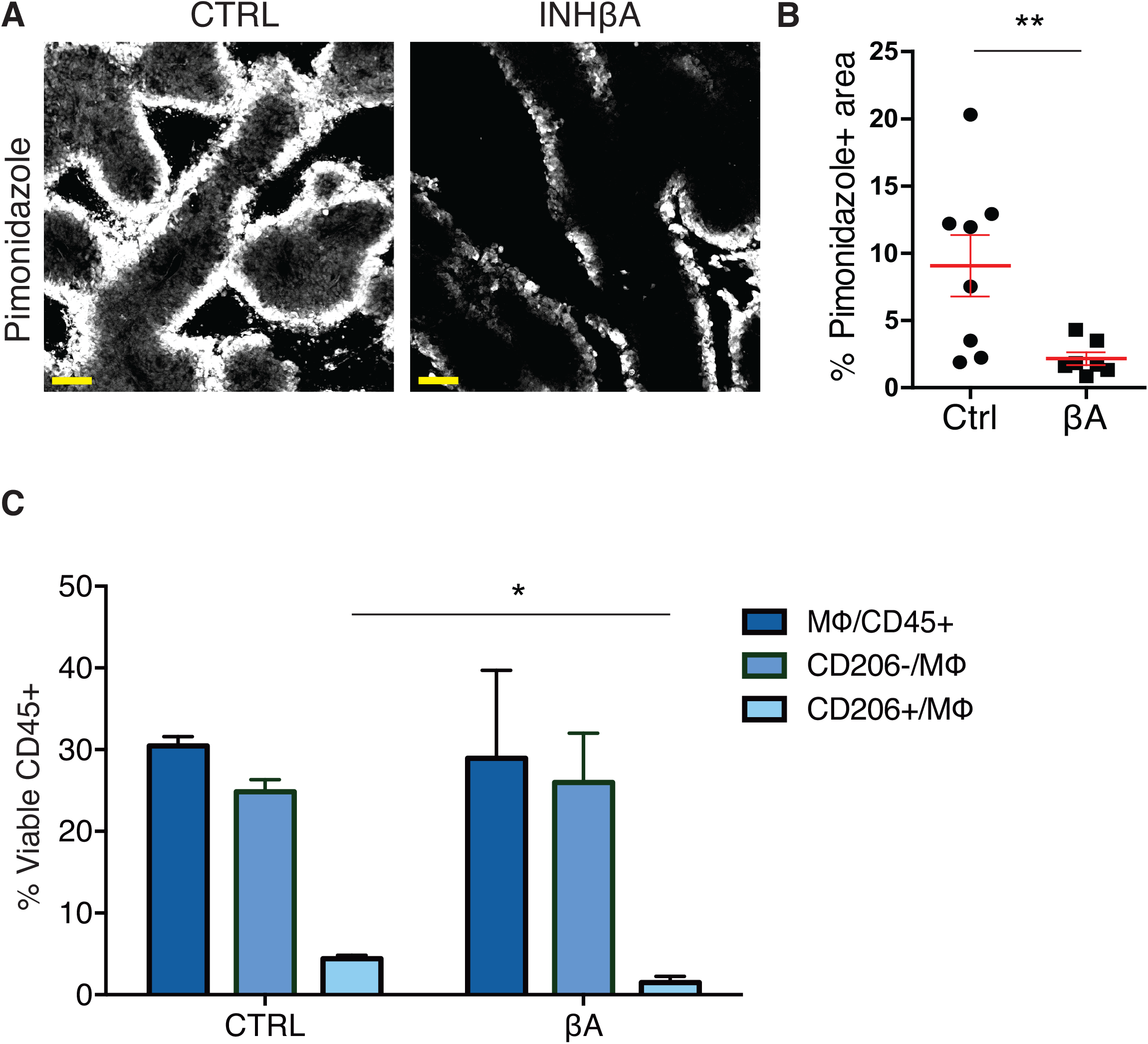
Hypoxic areas and tumor-infiltration by M2-like CD206-positive macrophages in B16F1:GFP melanoma. A) Pimonidazole immunofluorescent staining of hypoxic areas in 200 μm thick sections of CTRL or βA syngeneic intradermal B16F1:GFP melanoma grafts in wild-type C57BL/6 mice. B) Quantification of pimonidazole-positive area from entire sections. βA tumors had significantly decreased hypoxic areas (n=7) compared to CTRL (n=8), **p<0.01. C) Flow cytometric quantification of tumor-infiltrating F4/80+ macrophages. Total numbers (dark blue bars) and the frequencies of F4/80+CD11b+CD11c+CD206− M1-like macrophages (CD206−) were similar in CTRL and βA tumors. By contrast, F4/80+CD11b+CD11c+CD206+ M2-like macrophages were decreased in βA tumors (1.5%±0.7) compared to CTRL (4.4%±0.4), n=3 each, *p<0.05.

**Figure S9.**
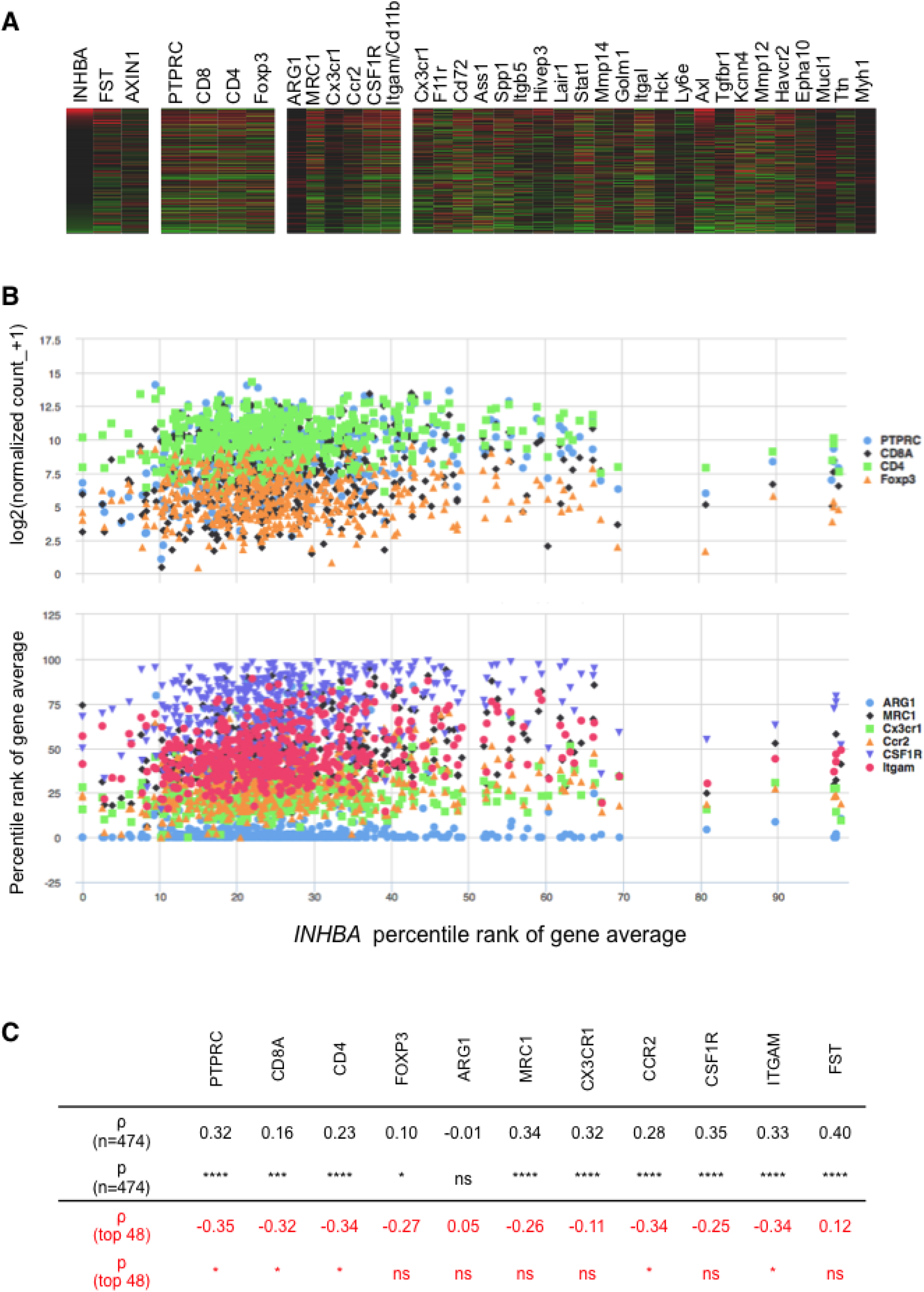
Expression of selected marker genes compared to *INHβA* mRNA levels in the TCGA Skin Cutaneous Melanoma dataset. A) Heatmaps of *INHβA* and *FST* expression compared to the Wnt target gene *AXIN1* (first panel from left), CD45 (PTPRC) and other markers of T lymphocyte subsets (second panel), myeloid markers (third panel) and of a set of genes that are regulated by Activin-A in monocyte/macrophages in mouse skin squamous cell carcinoma models (Antsiferova et al., 2016) (fourth panel). B) Expression of the indicated lymphocyte (top panel) or myeloid markers (bottom panel) plotted by the Xenabrowser against the percentile rank of *INHβA* expression. In the bottom panel, the y-axis shows percentile rank of gene expression instead of normalized log2 values in order to obtain better visual resolution of data points. C) Spearman (ρ) correlation between normalized mRNA levels (log2 values) of the indicated genes and *INHβA* mRNA expression. Note that in the 10% of tumors with the highest *INHβA* expression (n=48), 5 of 11 markers correlated negatively, whereas a significant positive correlation was observed across the entire dataset for 10 of them (n=474). *p<0.05, ***p<0.001, ****p<0.0001.

**Supplemental Table S1.**
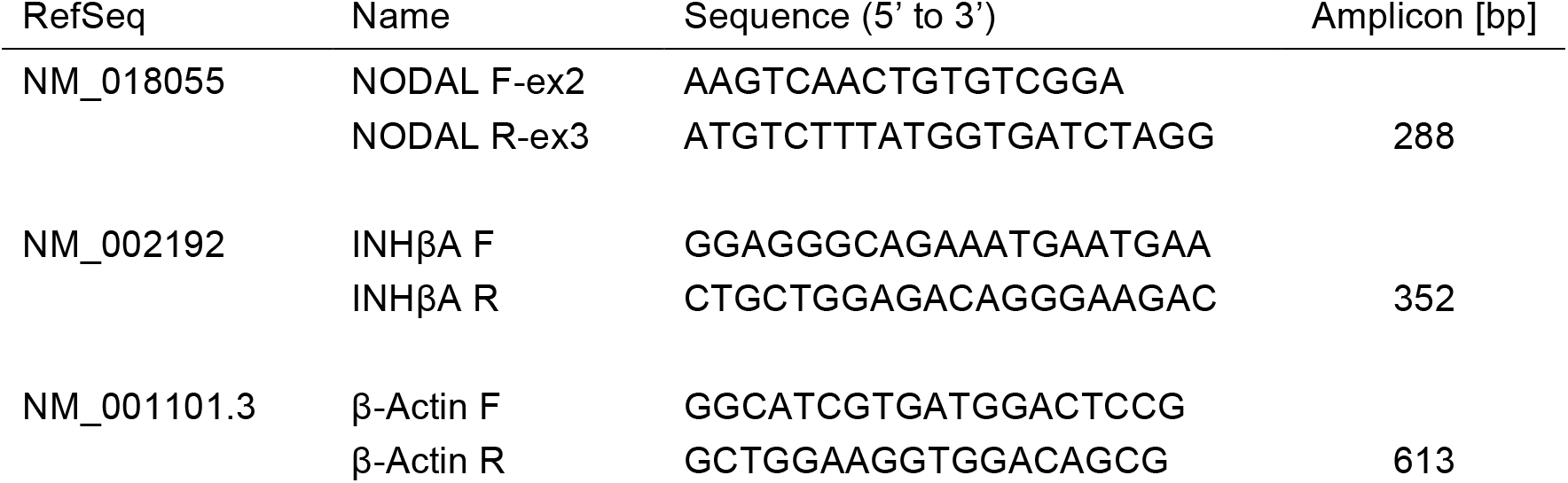
RT-PCR primers

